# Regulation of RNA methylation by therapy treatment, promotes tumor survival

**DOI:** 10.1101/2023.05.19.540602

**Authors:** Syed IA Bukhari, Samuel S Truesdell, Chandreyee Datta, Pritha Choudhury, Keith Q Wu, Jitendra Shrestha, Ruby Maharjan, Ethan Plotsker, Ramzi Elased, Sadia Laisa, Vijeta Bhambhani, Yue Lin, Johannes Kreuzer, Robert Morris, Siang-Boon Koh, Leif W. Ellisen, Wilhelm Haas, Amy Ly, Shobha Vasudevan

## Abstract

Our data previously revealed that chemosurviving cancer cells translate specific genes. Here, we find that the m6A-RNA-methyltransferase, METTL3, increases transiently in chemotherapy-treated breast cancer and leukemic cells in vitro and in vivo. Consistently, m6A increases on RNA from chemo-treated cells, and is needed for chemosurvival. This is regulated by eIF2α phosphorylation and mTOR inhibition upon therapy treatment. METTL3 mRNA purification reveals that eIF3 promotes METTL3 translation that is reduced by mutating a 5′UTR m6A-motif or depleting METTL3. METTL3 increase is transient after therapy treatment, as metabolic enzymes that control methylation and thus m6A levels on METTL3 RNA, are altered over time after therapy. Increased METTL3 reduces proliferation and anti-viral immune response genes, and enhances invasion genes, which promote tumor survival. Consistently, overriding phospho-eIF2α prevents METTL3 elevation, and reduces chemosurvival and immune-cell migration. These data reveal that therapy-induced stress signals transiently upregulate METTL3 translation, to alter gene expression for tumor survival.

**One sentence summary:** m6A enzyme translation upon therapy stress, promotes tumor survival

## Introduction

Previous findings revealed that cells that are resistant to chemotherapy (Lee et al., 2020), exhibit altered post-transcriptional gene expression. In such chemo-treated cancer cells, mTOR activity is transiently inhibited, and the integrated stress response (ISR), is activated (Gandin et al., 2016; Harvey et al., 2019; Jewer et al., 2020; Lee et al., 2020; Min and Spencer, 2019; Pakos-Zebrucka et al., 2016). These two stress response pathways suppress both rate-limiting steps of canonical cap dependent translation. mTOR inhibition blocks mRNA recruitment via the canonical cap complex, one of the two rate-limiting steps of canonical translation. ISR involves activation of eIF2α kinases (eIF2aks, PERK, GCN2, HRI, PKR) that phosphorylate the eIF2α subunit of the canonical tRNA recruiter eIF2 complex, leading to inhibition of the other rate-limiting step of conventional translation of recruitment of initiator tRNA (Costa-Mattioli and Walter, 2020; Sonenberg and Hinnebusch, 2009). These changes inhibit proliferation that is driven by canonical translation (Truitt and Ruggero, 2016), and permit specialized, post-transcriptional expression of distinct mRNAs to enable chemosurvival (Lee et al., 2020). How specific mRNAs are marked for selective post-transcriptional regulation in refractory cancer cells, to elicit chemosurvival, remains to be uncovered.

Modifications on mRNAs have been recently shown to cause their post-transcriptional regulation in distinct cellular conditions (Bringmann and Luhrmann, 1987; Jaffrey and Kharas, 2017; Liu et al., 2014; Meyer and Jaffrey, 2014; Roundtree et al.; Svitkin et al., 2017; Zhou et al., 2018). These alter structure, or mRNA and protein interactions, and recruit RNA binding proteins called readers that recognize the modification to cause post-transcriptional regulation of such mRNAs. Deregulation of the RNA methyltransferases (writers), their RNA binding protein effectors (readers) or their demethylases (erasers) have been implicated in various diseases including cancer (Bringmann and Luhrmann, 1987; Jaffrey and Kharas, 2017; Liu et al., 2014; Meyer and Jaffrey, 2014; Roundtree et al.; Svitkin et al., 2017; Zhou et al., 2018). Such modifications also mark cellular RNAs as self, to avoid triggering the cellular anti-viral immune response (Durbin et al., 2016; Karikó et al., 2005). The m6A RNA methyltransferase, METTL3, associates with its co-factor METTL14, to methylate the N6 position of Adenosine on mRNAs at RRACH motifs. METTL3 and its co-factor METTL14 have been implicated in the control of the stem cell state, in stress conditions and in cancers where their expression is deregulated, causing disease by altering m6A target RNA stability or translation changes (Jaffrey and Kharas, 2017; Meyer, 2019; Xiang et al., 2017). How m6A methyltransferases themselves may be regulated in chemosurviving cells remains to be fully explored.

In this study, we find that METTL3 and m6A are transiently upregulated in chemotherapy treated breast cancer and leukemic cells by therapy-induced stress signals that trigger eIF2α phosphorylation and mTOR inhibition. These stress signals decrease canonical translation in therapy-treated cells. We find that eIF2α phosphorylation enables non-canonical translation of METTL3. This is mediated by m6A modification in the 5′UTR of METTL3 mRNA that we find, binds the translation initiation factor, eIF3, that can bind m6A to promote translation (Hinnebusch, 2014; Meyer, 2019; Meyer et al., 2015a; Valášek et al., 2017). Importantly, the elevated METTL3 is needed for promoting chemosurvival and invasion, and for limiting the anti-viral immune response; these are reversed by overriding eIF2α phosphorylation to prevent METTL3 upregulation, or by depleting METTL3. We find that therapy treatment and METTL3 regulate PKR, one-carbon and other metabolism genes that alter the substrate for m6A methylation, S-adenosylmethionine (SAM), and the m6A demethylase activator, α-ketoglutarate (αKG) (Deng et al., 2018; Gerken et al., 2007; Li et al., 2017b), over time of therapy treatment; these impact methylation, to control METTL3 levels as feedback regulation. Together, our data reveal that METTL3 is regulated by therapy-induced stress response to promote m6A-mediated gene expression regulation for tumor survival.

## Results

### METTL3, and m6A on RNA, increase upon chemotherapy treatment in breast cancer and acute monocytic leukemic cell lines

To determine the mechanism of specific mRNA expression in chemosurviving cancer cells, we examined RNA binding regulators in MCF7 breast cancer cells, isolated after doxorubicin chemotherapy treatment. Tandem-mass-tag (TMT) spectrometry data of cells treated with or without doxorubicin, revealed an increase of RNA regulators including the m6A RNA methyltransferase, METTL3, by 3.5-fold (Table S1). We next conducted a time course with various concentrations of standard chemotherapies in breast cancer and acute monocytic leukemic cell lines. We find that METTL3 increases in doxorubicin-surviving MCF7 cells at multiple concentrations of doxorubicin treatment (250nM and 500nM) transiently over time within 18 h of treatment (Fig. 1A, S1A) or with other chemotherapies (Fig. S1B-C). This increase is not unique to MCF7 cells, as multiple concentrations of different therapies tested on triple negative breast cancer BT549 cells, or acute monocytic leukemic THP1 cells treated with a chemotherapeutic used in leukemia, Cytarabine (AraC), reveal similar transient increase in METTL3 (Fig. S1D-E). The upregulation observed is transient and is not observed at 48 h of chemotherapy treatment (Fig. S1F). These data suggest that METTL3 increases transiently, as a stress response.

**Fig 1.**
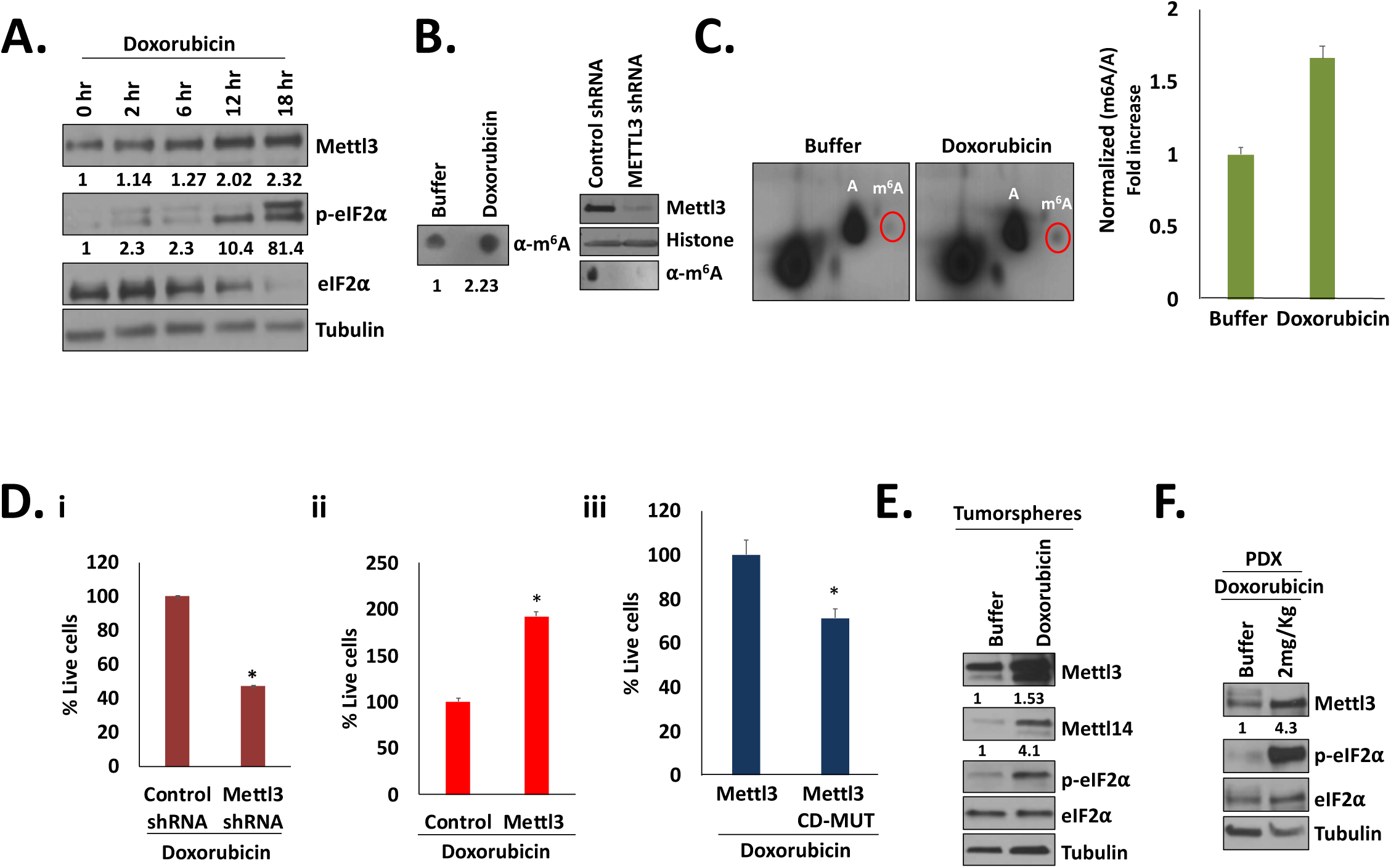
METTL3 and m6A modification on RNA are increased with chemotherapy doxorubicin treatment in breast cancer cells, and are needed for chemosurvival. **A.** Western blot analysis of METTL3 (identified from Table S1), and canonical translation marker: phospho-eIF2α and eIF2α, in MCF7 cells over time of treatment with doxorubicin (250nM) chemotherapy. **B.** Dot blots of rRNA and tRNA depleted cellular RNA from 500nM doxorubicin and control treated cells versus cells depleted of METTL3 with METTL3 shRNA, probed with m6A antibody. Western blot of samples with control shRNA or shMETTL3 shRNA. **C.** 2D-TLC of rRNA and tRNA depleted cellular RNA from 500nM doxorubicin treated and untreated MCF7 cells; m6A and A nucleosides are marked, and quantitated. **D.** Doxorubicin chemosurvival in MCF7 cells expressing **i.** control shRNA or METTL3 shRNA (500nM doxorubicin for 5 days) **ii.** wildtype METTL3 compared to a control GFP vector (500nM doxorubicin for 3 days), and **iii.** m6A catalytically defective METTL3 (CD-MUT) compared to wild type METTL3 (500nM doxorubicin for 3 days). **E.** Western blot show METTL3 and eIF2α phosphorylation increase in hormone positive breast cancer patient samples grown as 3D tumorspheres and treated with doxorubicin (250nM Doxorubicin for 18hrs), and in **F.** patient derived triple negative breast cancer xenografts (pdx) that were treated with 2mg/kg of doxorubicin in vivo for 1 hr. Tubulin and Histone serve as Western blot loading controls. Data are average of 3 replicates +/−SEM. See also Fig. S1, Table S1.

To test whether this leads to more m6A modification on RNA, we first tested cellular RNA after depletion of ribosomal RNA and of RNA less than 200nt (to avoid tRNAs), from doxorubicin-treated cells compared to RNA from untreated cells, by dot blot analyses. Consistent with the increased METTL3 levels (Fig. 1A, S1A-E), we find that doxorubicin-treated cells show more m6A-modified RNA compared to untreated cells (Fig. 1B). To verify these results, we next created a cell line with a constitutively expressed shRNA that efficiently depletes METTL3, and performed dot blot analysis. Consistent with decreased METTL3 upon knockdown, the dot blot signal is depleted when METTL3 is knocked down (Fig. 1B). Second, we tested RNA from doxorubicin-treated cells for m6A by two-dimensional-thin layer chromatography separation (2D-TLC) (Grosjean et al., 2004). We find that the m6A signal on 2D-TLC increases in ribosomal RNA-depleted, poly(A)-selected RNAs from doxorubicin-treated cells compared to untreated cells (Fig. 1C); this signal is reduced in RNA from shMETTL3-expressing cells compared to control shRNA cells (Fig. S1G), verifying the m6A signal. These data indicate that the elevated METTL3 enhances m6A on RNA in doxorubicin-treated cells.

### METTL3 is required for chemosurvival and increases with doxorubicin treatment in primary breast tumorspheres and in patient-derived xenografts

We next tested whether the increase of METTL3 and m6A in chemo-treated cells, is needed for chemosurvival. We depleted METTL3 using shRNA, or overexpressed METTL3 compared to a catalytically defective METTL3 (CD-M3) mutant (Lin et al., 2016), followed by doxorubicin treatment to test these cells for chemosurvival. We find that compared to a control shRNA, stable cells that efficiently deplete METTL3 (Fig. 1B), showed decreased chemosurvival (Fig. 1D). Correspondingly, overexpression of METTL3 showed increased chemoresistance while overexpression of CD-M3 catalytic defective mutant did not (Fig. 1D). These data suggest that the upregulation of METTL3; and thereby, of m6A on mRNA in chemotherapy-treated cells— that can alter gene expression—contributes to chemosurvival of such cells. Consistently, we find that METTL3 is elevated in hormone positive breast cancer patient samples grown as tumorspheres and treated with doxorubicin (Fig. 1E). Increase of METTL14 is also observed with doxorubicin where METTL3 is increased (Fig. 1E), consistent with the known co-regulated levels of these co-factors (Vu et al., 2017). Furthermore, we find that METTL3 is elevated in vivo, in doxorubicin-treated, triple negative breast cancer patient-derived xenografts (pdx, Fig. 1F). These data suggest that the increase of METTL3, observed in chemotherapy treated cell lines, is not an artifact and can be used to study its regulation, as METTL3 increases in vivo in chemosurviving, primary breast cancer.

### METTL3 and m6A increase with induction of eIF2**α** phosphorylation and mTOR inhibition

Multiple DNA damage drugs such as doxorubicin, AraC, gemcitabine, and mitotic inhibitors such as paclitaxel enhance METTL3 (Fig. S1B-C, E), suggesting that METTL3 increases in response to stress signals. Chemosurviving cells exhibit stress signaling, including mTOR inhibition, and transient activation of eIF2α kinases that phosphorylate and inhibit eIF2α (Gandin et al., 2016; Harvey et al., 2019; Jewer et al., 2020; Lee et al., 2020). Consistently, we find that doxorubicin treatment in breast cancer cells and AraC treatment in THP1 cells, promote eIF2α phosphorylation that correlated with METTL3 increase in Fig 1A and S1A-E. This was also observed in patient samples and in vivo (Fig. 1E-F), suggesting that eIF2α phosphorylation may be important for METTL3 increase. Consistently, if we induce eIF2α phosphorylation by double-strand RNA mimic, poly I:C, or by thapsigargin (Berlanga et al., 2006; Costa-Mattioli and Walter, 2020; Harding et al., 2003; Muaddi et al., 2010; Sidrauski et al., 2013; Wek, 2018), we find that METTL3 and METTL14 are increased (Fig. 2A, S2A). In accord, we find increased m6A signal on dot blots of ribosomal RNA-depleted poly (A) RNAs derived from poly I:C treated cells (Fig. 2B) where METTL3 increases. These data indicate upregulation of METTL3 and m6A modification of RNAs, concurrent with induction of eIF2α phosphorylation.

**Fig. 2.**
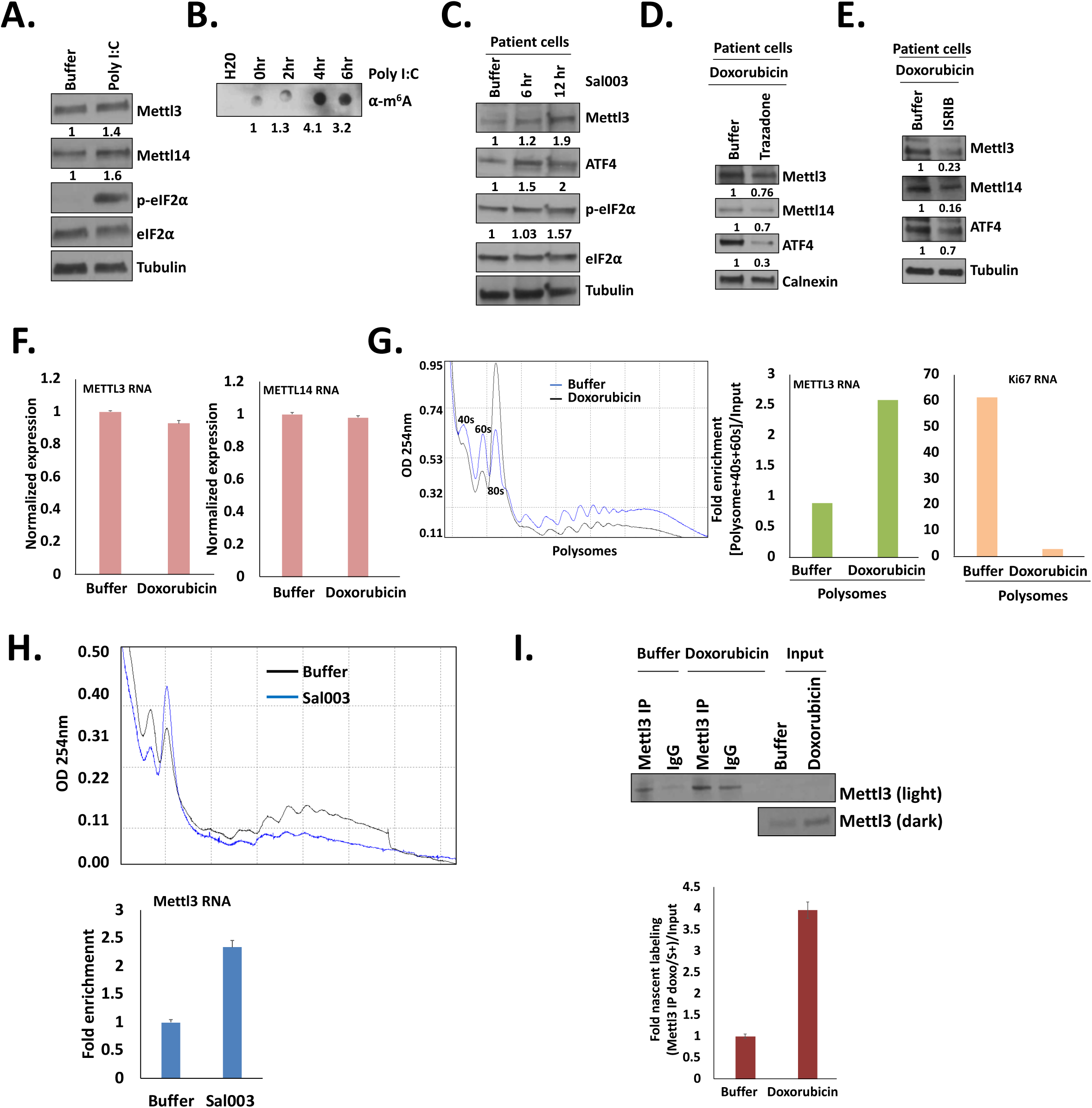
eIF2α phosphorylation promotes METTL3 translation. Western blots of MCF7 cells treated with **A.** poly I:C to induce eIF2α phosphorylation (2ug/ml). **B.** m6A dot blot analysis with poly I:C treatment. **C.** Treatment of hormone positive patient sample BT29 and BT30 cells with Sal003 (10uM) phosphatase inhibitor to retain eIF2α phosphorylation. **D.** Western blot of METTL3, METTL14 and ATF4 levels that do not increase when 500nM doxorubicin-treated patient sample cells that are also co-treated with 5uM Trazodone, or with **E.** with 1uM ISRIB, to override the effect of eIF2α phosphorylation. **F.** qPCR analysis of METTL3 and METTL14 RNA levels in 500nM doxorubicin (1day) treated MCF7 cells, compared to untreated cells. Polysome fractionation followed by qPCR analysis of polysome fractions **G.** for METTL3 mRNA and KI67 mRNA in 500nM doxorubicin (1day) treated MCF7 cells compared to untreated cells. **H.** for METTL3 mRNA (below) in Sal003 (10uM 1 day) treated MCF7 cells compared to untreated cells. **I.** Nascent amino acid labeling and METTL3 immunoprecipitation (Western blot of METTL3 with dark exposure of 1/10 Inputs) from 500nM doxorubicin (1day) treated versus untreated MCF7 cells to detect translation levels (quantitation below) of newly labeled immunoprecipitated METTL3 in these cells. Tubulin and Calnexin serve as Western blot loading controls. Data are average of 3 replicates +/−SEM. See also Fig. S2.

Chemotherapy-treated cells also show mTOR and PI3K inhibition that can activate PKR, and lead to eIF2α phosphorylation (Gandin et al., 2016; Harvey et al., 2019; Jewer et al., 2020; Lee et al., 2020). METTL3 has been shown to be increased upon mTOR inhibition (Zhou et al., 2018). Therefore, we treated MCF7 cells with Torin1 that blocks both mTORC1 and mTORC2 (Thoreen et al., 2012), to test whether METTL3 would increase along with enhanced eIF2α phosphorylation. As shown in Fig. S2B, we find that Torin1 treatment increases METTL3 along with eIF2α phosphorylation; 4EBP dephosphorylation marked the efficacy of Torin 1 inhibition of mTOR. In contrast, treatment with mTOR activator, MHY1485 (Choi et al., 2012), exhibits decreased METTL3 and eIF2α phosphorylation. METTL3 enhancement was also observed with Palomid 529 (Xue et al., 2008), an mTORC1 and mTORC2 inhibitor (Fig. S2C) and with BEZ 235, a dual inhibitor (Fig. S2D) of mTOR and PI3K that prevents reactivation of mTOR (Muranen et al., 2012). Elevated METTL3 and METTL14, with concordantly increased m6A on RNA (Fig. S2E), are also observed in Torin1-treated MCF7 tumorspheres that mimic in vivo tumors (Muranen et al., 2012). Thus, we find that mTOR inhibition, shows elevation of METTL3 along with eIF2α phosphorylation (Fig. S2B-E).

### Elevation of METTL3 and m6A, requires eIF2**α** phosphorylation

The common feature in all these conditions (Fig. 1, S1, 2A-B, S2A-E) that promote METTL3 levels is the concurrent increase in eIF2α phosphorylation. To test the need for eIF2α phosphorylation for METTL3 elevation, we used an inhibitor of eIF2α phosphatase, Sal003 that would retain eIF2α phosphorylation (Robert et al., 2006; Zismanov et al., 2016). Sal003 treatment in MCF7 or patient-derived hormone positive breast cancer cells, increased phospho-eIF2α, and ATF4, an ISR response gene that is non-canonically translated due to increased eIF2α phosphorylation (Hinnebusch, 2014; Muaddi et al., 2010; Pakos-Zebrucka et al., 2016; Wek, 2018); consistently, we find that Sal003 increased METTL3 (Fig. 2C). Concordantly, 2D-TLC analysis shows that Sal003 treatment elevated m6A on RNA, compared to control buffer treatment of MCF7 cells (Fig. S2G). When we tested overexpression of 4EBP mutants or eIF2α phospho mimetics, transfection stress caused transient eIF2α phosphorylation, preventing clear analysis (data not shown). These data suggest that induction of eIF2α phosphorylation may promote METTL3 levels.

eIF2α phosphorylation can be bypassed pharmacologically: by ISRIB, which modifies the guanine nucleotide exchange factor for eIF2α, eIF2B, to override the effect of eIF2α phosphorylation and enable ternary complex formation (Costa-Mattioli and Walter, 2020; Sidrauski et al., 2013; Zyryanova et al., 2018), or by trazodone that is thought to override eIF2α phosphorylation at the ternary complex stage (Harvey et al., 2019). We co-treated cells with doxorubicin with or without these ISR inhibitors, and as a control, checked the levels of ATF4 that is not increased in the presence of ISRIB or trazodone (Costa-Mattioli and Walter, 2020; Lebeaupin et al., 2020; Sidrauski et al., 2013; Vasudevan et al., 2020; Zyryanova et al., 2018). As shown in Fig. 2D and 2E, we find that doxorubicin-treated patient sample derived breast cancer cells that are also treated with trazodone or ISRIB to override phospho-eIF2α, do not show increase of ATF4, and also do not show increase of METTL3 levels, compared to buffer-treated cells. Torin 1-treated MCF7 cells also increase METTL3 co-incident with eIF2α phosphorylation (Fig. S2B); therefore, we tested whether this increase of METTL3 with Torin 1, needed the co-incident phosphorylation of eIF2α to elevate METTL3, by overriding phospho-eIF2α with ISRIB. In accord with the need for eIF2α phosphorylation, we find that co-treatment with Torin 1 and ISRIB reduced METTL3 (Fig. S2F). These data suggest that chemotherapy induces eIF2α phosphorylation to increase METTL3.

### METTL3 upregulation in doxorubicin-treated cells, is due to enhanced translation

As eIF2α phosphorylation suppresses canonical translation and permits non-canonical translation of specific mRNAs, these data suggested that METTL3 may be increased via non-canonical translation in therapy-treated cells. Consistently, no increase of METTL3 and METTL14 were seen at the RNA levels by qPCR analysis in doxorubicin-treated cells (Fig. 2F), further supporting that METTL3 and METTL14 are increased at the translation or post-translation levels. To verify that the increase in METTL3 was at the translation level, first, polysome analysis was conducted. We find that METTL3 mRNA increased on polysomes compared to monosomes, in doxorubicin-treated cells compared to buffer treated cells; in contrast, the cell cycle marker KI67 mRNA that is not translated in these arrested conditions decreased on polysomes (Fig. 2G). Second, to show that the translation effect on METTL3 was induced by eIF2α phosphorylation, polysome analysis was performed with the eIF2α phosphatase inhibitor, Sal003. As shown in Fig. 2H, METTL3 mRNA increased on polysomes compared to monosomes in Sal003-treated cells compared to buffer-treated cells, showing that METTL3 mRNA is translated in conditions of eIF2α phosphorylation. Third, we used Y10B antibody that was previously demonstrated to bind 5.8S rRNA in assembled ribosomes and has been used to show ribosome association (Lerner et al., 1981); consistent with METTL3 translation and thus ribosome association, Y10B antibody immunoprecipitated METTL3 and METTL14 mRNAs in doxorubicin and Torin 1 treated cells (Fig. S2H). Fourth, to ensure that the METTL3 mRNA polysome association is not indirect and METTL3 increase is due to enhanced translation activity, we used L-homopropargylglycine (HPG), an amino acid analog of methionine containing an alkyne moiety that can be biotinylated by click-it chemistry (ThermoFisher), for labeling nascently-translated proteins, followed by SDS-PAGE and HRP-streptavidin Western analysis. We find that immunoprecipitation of METTL3 (Fig. 2I, S2I), revealed an increase of HPG-labeled METTL3 in doxorubicin-treated cells compared to untreated cells, indicating increased translation of METTL3 protein in doxorubicin-treated cells. These data indicate that METTL3 mRNA is translationally upregulated in doxorubicin-resistant cells upon eIF2α phosphorylation.

### eIF3 binds METTL3 mRNA and is needed for METTL3 translation upregulation

To identify proteins that associate with METTL3 mRNA to promote its non-canonical translation, we used a biotin tag antisense (Borah et al., 2011) to METTL3 and METTL14 mRNAs. The antisense are against the shRNA target sites, as these sites are verified to be available for base pairing with the antisense, compared to a scrambled control antisense. We combined this with in vivo formaldehyde (for complexes) and UV (for direct interactions) crosslinking of doxorubicin-treated and untreated MCF7 cells, to ensure purification of true in vivo frozen complexes. We analyzed the antisense purification of METTL3 mRNA for association with RNA binding proteins and translation factors that are specifically increased in chemo-treated cells, identified by TMT spectrometry (Fig. 3A, Table S1, S2A). Purification of METTL3 mRNA, from in vivo formaldehyde crosslinked extracts from doxorubicin treated cells (Fig. 3A), revealed that METTL3 mRNA, but not a control antisense, associates with eIF3 complex translation factors, and other proteins (Table S2A) that are also increased in doxorubicin-treated cells (Table S1). To look for direct binding, we examined extracts that were UV crosslinked. We find that eIF3 co-purifies with METTL3 and METTL14 mRNAs but not with a control scrambled antisense (Fig. 3A). METTL14 mRNA co-regulation with METTL3 levels has been noted earlier (Vu et al., 2017). We also conducted m6A antibody immunoprecipitation from formaldehyde crosslinked doxorubicin-treated and untreated MCF7 cells. These m6A antibody immunoprecipitates may inform on the complexes associated with RNAs that are m6A modified. Consistently, these immunoprecipitates reveal RNA processing factors as well as eIF3 (Table S2B) that was also observed by antisense purification (Fig. 3A, S3A). In accord, UV crosslinking and eIF3 immunoprecipitation reveals associated METTL3 mRNA (Fig. S3B). EIF3 is a multiprotein complex that can recruit the ribosome for canonical and non-canonical translation initiation (Hinnebusch, 2014; Valášek et al., 2017). EIF3 subunit components such as those identified (eIF3 d, e, g, m, a, Fig. 3A, S3A-B, Table S2A) can initiate non-canonical translation, including by recognizing m6A sites in the 5′UTRs of specific mRNAs (Choe et al., 2018; Meyer et al., 2015b; Sun et al., 2013). Consistently, we find that overexpression of eIF3d promoted METTL3 levels and accordingly, enhanced chemosurvival (Fig. 3B-C), while depletion of eIF3a reduced METTL3 levels and chemosurvival (Fig. 3D-E). These data suggest that eIF3 associates with METTL3 mRNA to promote its non-canonical translation.

**Fig 3.**
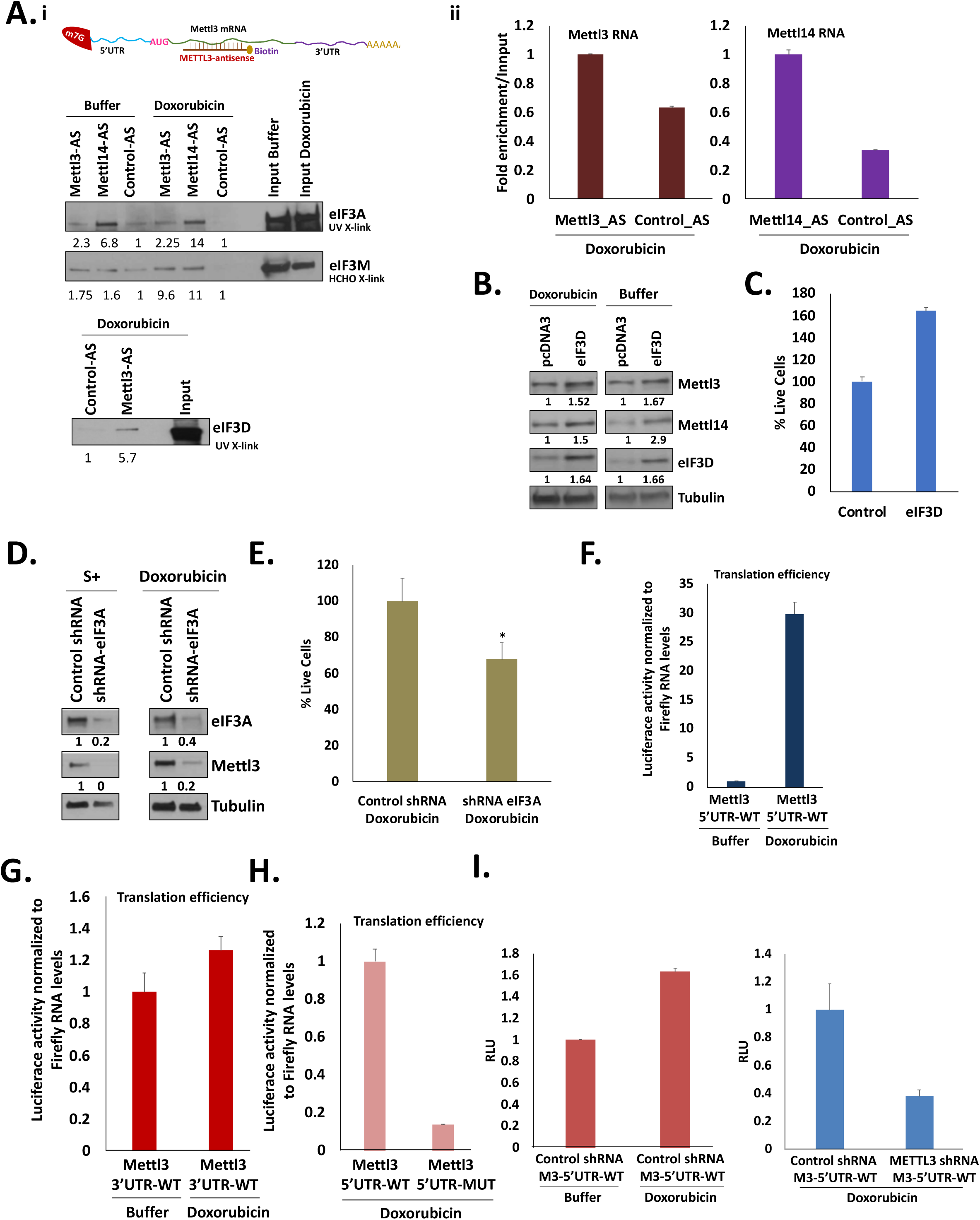
EIF3 promotes METTL3 translation in doxorubicin-treated MCF7 cells. **A. i.** Western blot of eIF3a, eIF3m, and eIF3d in eluates after streptavidin affinity purification by biotin-tagged antisense of METTL3 and METTL14 mRNAs compared to a scrambled control antisense, from in vivo UV and formaldehyde (HCHO) crosslinked, 250 nM and 500 nM doxorubicin (12 h and 1 day) treated or untreated MCF7 cells (scheme shown above, data in Table S2A). **ii.** qPCR of METTL3 and METTL14 RNAs co-purified by antisense compared to control antisense in doxorubicin treated cells. **B.** Western blot of METTL3 in MCF7 cells with eIF3d overexpression compared to control, which are 500nM doxorubicin (1day) treated or untreated. **C.** Chemotherapy survival cell counts of eIF3d overexpressing MCF7 cells compared to control cells, treated with 500nM doxorubicin for 20 h. **D.** Western blot of eIF3a depletion compared to control vector, transfected in MCF7 cells, followed by 500nM doxorubicin treatment to test **E.** chemotherapy survival by percentage surviving cell count after 3 days. Translation efficiency of Firefly luciferase reporter bearing **F.** the 5′UTR of METTL3 normalized to co-transfected Renilla in untreated and 500nM doxorubicin (1 day) treated MCF7 cells, normalized for reporter RNA levels, or **G.** bearing the 3′UTR of METTL3. **H.** Translation efficiency of Firefly luciferase reporter bearing the 5′UTR of METTL3 compared to an identical reporter bearing mutated GAC (m6A modification motif mutated to GUC) site upstream of the reporter start site in the 5′UTR of METTL3, normalized to co-transfected Renilla and reporter RNA levels, in 500nM doxorubicin (1 day) treated MCF7 cells. **I.** Translation of Firefly luciferase reporter bearing the 5′UTR of METTL3 in METTL3 shRNA MCF7 cells compared to control shRNA cells, with or without 500nM doxorubicin treatment for 1 day. Tubulin is the loading control for Western blots. Data are average of 3 replicates +/−SEM. See also Fig. S3, Table S1, S2A-B.

### METTL3 5**′**UTR promotes reporter translation in response to doxorubicin treatment

To test for cis-acting elements on METTL3 mRNA, which may direct its translation upregulation in doxorubicin-treated MCF7 cells, we constructed Luciferase reporters bearing METTL3 mRNA 5′UTR and 3′UTR, or their reverse sequences or vector sequence as controls. We transfected these reporters in untreated and doxorubicin-treated MCF7 cells, and normalized Luciferase activity for luciferase RNA levels by qPCR, to calculate translation efficiency. We find that METTL3 5′UTR reporter promoted translation efficiency over 3-fold, in doxorubicin-treated cells, compared to untreated cells, mimicking the endogenous METTL3 (Fig. 3F). The 3′UTR reporter did not upregulate translation in doxorubicin-treated cells compared to untreated cells (Fig. 3G), indicating that the 3′UTR was not involved in translation upregulation upon doxorubicin treatment. These data indicate that METTL3 5′UTR promotes translation upon doxorubicin treatment.

### METTL3, and the m6A site in METTL3 5**′**UTR, are needed to promote METTL3 5**′**UTR reporter translation

While m6A modification causes RNA downregulation, it can also promote non-canonical translation in the 5′UTR via eIF3 recruitment to the m6A-modified GAC motif (Meyer et al., 2015b). We identified that the METTL3 mRNA binds eIF3 (Fig. 3A, S3) and needs the 5′UTR for translation upregulation in doxorubicin-treated cells (Fig. 3F). These data suggest that m6A-mediated translation may be happening on METTL3 mRNA that harbors RRACH (GAC) motifs upstream of the ATG. Therefore, we constructed reporters with METTL3 5′UTR with the GAC site mutated. We find that while the 5′UTR reporter increased in doxorubicin-treated cells compared to untreated cells, the GAC mutant reporter showed no significant increase (Fig. 3H). This suggested that the presence of METTL3 levels or m6A activity to modify the GAC site, may regulate its own 5′UTR reporter translation. To verify the role of m6A on its 5′UTR reporter translation, we expressed METTL3 5′UTR luciferase reporter in MCF7 cells with and without METTL3 depletion. Consistently, depletion of METTL3 decreased translation of the 5′UTR reporter, compared to control shRNA in doxorubicin-treated cells (Fig. 3I). These data suggest that METTL3 may regulate its own levels via m6A modification of its 5′UTR to promote its non-canonical translation in doxorubicin treated cells.

### METTL3 downregulates proliferation genes

To identify METTL3 targets in therapy-treated breast cancer cells, we conducted global profiling analysis at multiple levels. We identified m6A-modified RNAs by co-immunoprecipitation with antibody against m6A (meRIP), from doxorubicin-treated compared to untreated MCF7 cells, to identify associated target RNAs that bear m6A marks (Table S3). Consistent with our data (Fig. 3F-I), we find that METTL3 mRNA is also immunoprecipitated. We also profiled shMETTL3 cells compared to control shRNA vector cells upon doxorubicin treatment at the transcriptome (Table S3) and proteome (Table S4) levels to identify genes that are regulated upon METTL3 depletion in chemo-treated cells (Table S3-4), and selected those that were also m6A targets (1.5 fold and greater) from the meRIP in doxorubicin treated cells but not in untreated cells (Fig. 4Ai, Table S5 for commonly regulated genes from Table S3-S4). METTL14 mRNA co-regulation with METTL3 levels has been noted earlier (Vu et al., 2017), and is observed in our proteomic data (Table S4). We find that predominantly, cell cycle genes are decreased by METTL3 at the RNA levels, and are the top category associated with m6A antibody (Fig 4Aii); consistently, these increase upon METTL3 depletion in MCF7 cells (Fig 4Aiii). As depletion of METTL3 increases cell cycle mRNAs (Fig. 4Aiii), it would render the cells vulnerable to chemotherapy due to increased cell cycle and replication; consistently, we find that depletion of METTL3 or eIF3 that is required for increased METTL3, reduced chemosurvival (Fig. 1D, 3E). In contrast, METTL3 overexpression (Fig. 1D) or overexpression of eIF3 that increases METTL3 (Fig. 3D), promoted chemosurvival, while expression of a catalytically defective mutant of METTL3 (CD-M3) reduced chemosurvival (Fig. 1D). These data suggest that the elevated METTL3 in chemo-treated cells, downregulates proliferation mRNAs to suppress chemosensitivity.

**Fig. 4.**
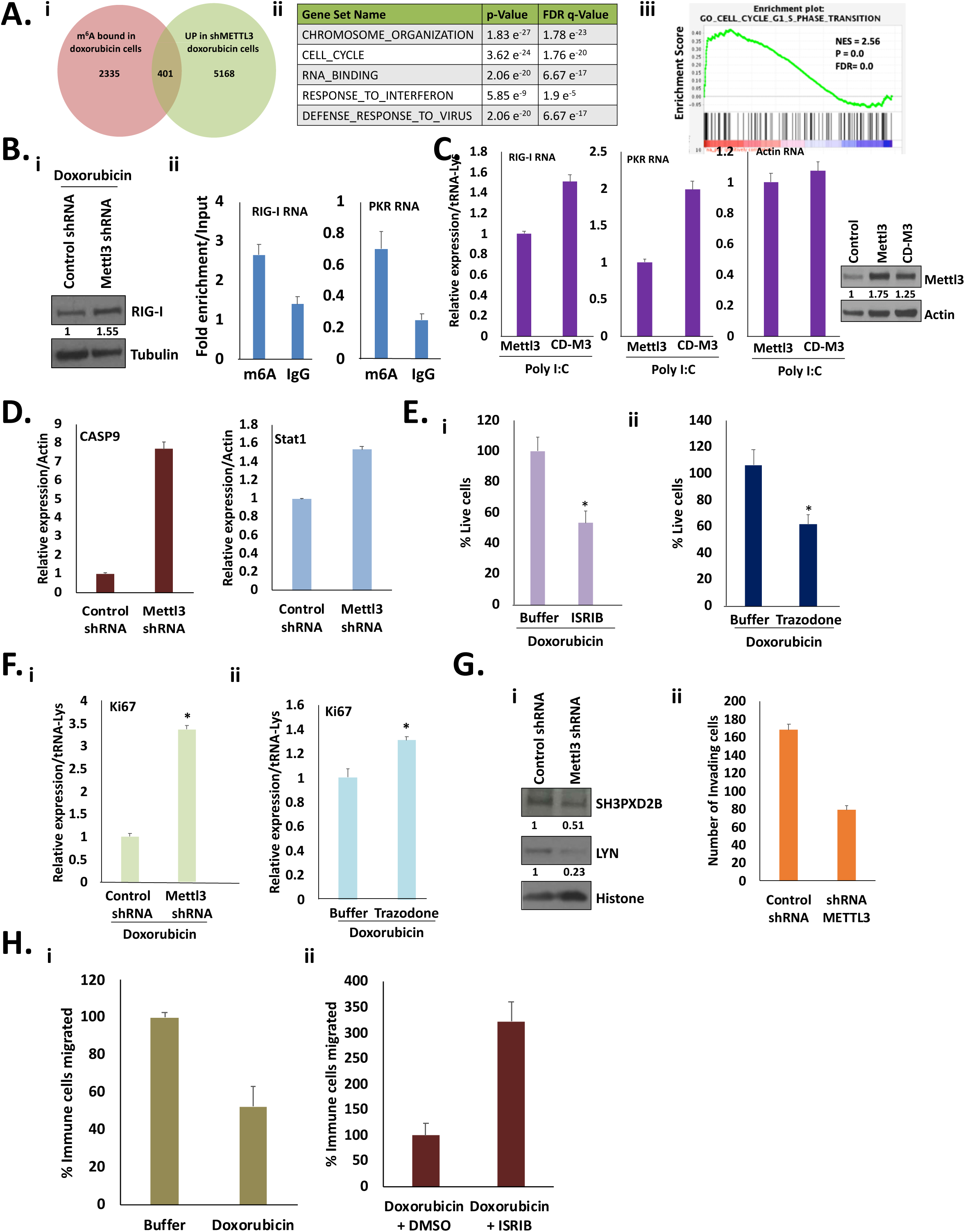
METTL3 suppresses proliferation and anti-viral immune response, while promoting invasion genes, causing chemosurvival and limiting immune cell migration. **A. i.** m6A IP RNAs from 500nM doxorubicin treated (1 day) cells but not in untreated cells (2335, Table S3) were compared with genes upregulated in shMETTL3 cells versus control shRNA cells (5168, Table S3 for RNA levels, Table S4 for protein levels) to identify RNAs that are m6A marked and downregulated by METTL3 (401, Venn Diagram, Table S5 for commonly identified genes from Table S3-S4). Upregulated genes were also compared but were fewer. **ii.** RNAs that are m6A modified: cell cycle RNAs (Ki67, PLK1), and anti-viral (DDX58 or RIG-I, PKR) RNAs but not control RNAs such as Actin and tRNA-lys are associated with m6A antibody and are increased upon METTL3 depletion in shMETTL3 MCF7 cells at the protein and RNA levels by TMT spectrometry and microarray respectively. **iii.** GSEA analysis of m6A-associated RNAs, reveals enrichment of cell cycle genes that are significantly (>=1.5 fold) upregulated upon METTL3 depletion. **B. i.** Western blot analysis of RIG-I anti-viral protein in shMETTL3 compared to control shRNA expressing MCF7 cells treated with 500nM doxorubicin for 1 day. **ii.** M6A antibody immunoprecipitation of RIG-I and PKR anti-viral gene mRNAs compared to IgG control. **C.** Anti-viral response to treatment with poly I: C (2ug/ml for 18 hrs) was tested by qPCR levels of RIG-I and PKR RNAs compared to control Actin mRNA by qPCR. This was tested in METTL3 overexpressing MCF7 cells compared to catalytic mutant METTL3 CD-M3 overexpressing cells. Shown Western blot of METTL3 levels in MCF7 cells with METTL3 and CD-M3 overexpression, compared to control vector. **D.** CASP9 and STAT1 downstream targets of RIG-I, are increased in METTL3 depleted MCF7 cells, as observed by qPCR analysis normalized for Actin mRNA. **E.** Inhibition of doxorubicin chemosurvival in MCF7 cells, with small molecules that override eIF2α phosphorylation (trazodone, ISRIB), compared to buffer treated control. 500nM Doxorubicin-treated MCF7 cells were co-treated with **i.** 1uM of ISRIB, **ii.** 5uM of Trazodone or DMSO buffer. Western blot analyses of METTL3 and METTL14 levels and ATF4 levels are shown with these (Trazodone or ISRIB combined with doxorubicin) treatments in Fig. 2D-E compared to buffer treated control cells. **F.** Ki67 cell cycle gene increases in **i.** Doxorubicin-treated shMETTL3 MCF7 cells compared to control shRNA cells, and in **ii.** Doxorubicin-treated MCF7 cells that are co-treated with 5uM of Trazodone, compared to DMSO buffer. **G. i.** Western blot showing decrease of invasion genes upon METTL3 depletion in shMETTL3 MCF7 cells compared to control shRNA cells. **ii.** Invasion assay showing decrease in invading cells upon METTL3 knockdown compared to control shRNA MCF7 cells. **H. i.** Bar graph shows fewer CD14+ monocytes migrate towards doxorubicin-treated MCF7 cells where METLL3 levels are increased compared to untreated MCF7 cells. **ii.** Bar graph shows more CD14+ monocytes migrate towards doxorubicin-treated MCF7 cells that were co-treated with ISRIB that prevents increase of METTL3 levels. Data are average of 3 replicates +/−SEM. Actin and Tubulin are loading controls for Western blots. See also Fig. S4, Table S3-5.

### METTL3 downregulates anti-viral immune response genes

A second class of genes affected by METTL3 depletion and m6A antibody are anti-viral immune response genes (Fig 4Aii, Table S3-S5), such as PKR and DDX58/RIG-I. Such genes trigger innate immune response that can cause immune cell migration and a pro-inflammatory response that promotes cell death (Bennett et al., 2012; Besch et al., 2009; Darini et al., 2019; Elion et al., 2018; Peidis et al., 2011). We find that RIG-I and PKR that recognize unmodified RNA, are increased upon METTL3 depletion (Fig. 4B). As m6A methylation on RNA causes RNA to be recognized as self and does not trigger the anti-viral response (Durbin et al., 2016; Karikó et al., 2005), this would complement the loss of m6A in shMETTL3 cells to promote the RIG-I response. If METTL3 depletion increases the anti-viral response, then overexpression of METTL3 would reduce anti-viral response. This would also be consistent and complementary to the role of m6A in masking RNAs to preclude the anti-viral immune response (Durbin et al., 2016; Karikó et al., 2005). We tested this by overexpressing METTL3 or the catalytically defective mutant CD-M3 (Lin et al., 2016), followed by poly I:C treatment to test for anti-viral response gene upregulation by qPCR. We found that compared to the catalytically defective mutant CD-M3 expressing cells, cells overexpressing METTL3 reduced the anti-viral response mRNAs of RIG-I and PKR (Fig. 4C). The RIG-I anti-viral immune response can contribute to cell death (Besch et al., 2009; Boelens et al., 2014; Li et al., 2017a; Majzoub et al., 2019) of METTL3 depleted cells. In accord, we find that downstream targets of RIG-I that mediate cell death and anti-viral innate immune response (Elion et al., 2018), CASP9 (Tang et al., 2019) and STAT1 (Jiang et al., 2011; Yoneyama et al., 2004), are increased in METTL3 depleted cells (Fig. 4D); increase of such cell death regulators upon RIG-I elevation due to loss of m6A regulation, is consistent with the increased cell death seen in METTL3 depleted cells (Fig. 1D). These data suggest that therapy stress signals increase METTL3 to suppress the anti-viral response that can lead to innate immune response and cell death. With decreased levels and activation of the anti-viral gene PKR, due to elevated METTL3 upon doxorubicin treatment (Fig. 4A-C), eIF2α phosphorylation would decline, preventing non-canonical translation of METTL3 itself as a feedback regulation loop. Consistently, we see reduced eIF2α phosphorylation, and METTL3 over later time points of doxorubicin treatment (Fig. 1A, S1A-F). This transient regulation of METTL3, and PKR could allow for ISR-induced METTL3-mediated gene expression regulation for survival, while preventing cell death effects of prolonged levels of PKR and phospho-eIF2α (Bennett et al., 2012; Darini et al., 2019; Peidis et al., 2011).

### Overriding EIF2**α** phosphorylation reduces chemosurvival

Our data suggest that stress signals in chemotherapy-treated cells cause eIF2α phosphorylation that enables METTL3 upregulation, which alters gene expression to promote chemosurvival by suppressing cell cycle and cell death regulators. Therefore, pharmacological inhibitors that override eIF2α phosphorylation (ISRIB, trazodone), could alleviate chemosurvival. Consistent with our results with METTL3 depleted cells, doxorubicin-treated cells co-treated with trazodone or ISRIB that prevent the increase of METTL3 in Fig. 2E-F, correspondingly show significant decrease in chemosurvival and consistently increased cell cycle genes (Fig. 4E-F). Synergistic decrease in chemosurvival was not observed upon METTL3 depletion and trazodone, indicating that the eIF2α pathway mediates chemosurvival at least in part through METTL3 (Fig. S4A). This is consistent with the increase of anti-viral and cell cycle genes upregulated in shMETTL3 cells that correlate with decreased chemosurvival. These data suggest that chemosurvival can be reduced by suppressing the effect of the eIF2α phosphorylation pathway. This reduces METTL3 and thus prevents m6A regulation of mRNAs that need to be controlled to support chemosurvival.

### Doxorubicin-induced METTL3 promotes expression of invasion genes

m6A can also promote non-canonical translation (Choe et al., 2018; Meyer et al., 2015b; Zhou et al., 2018). Therefore, we examined our meRIP datasets compared to Mettl3 depletion RNA profiles and proteomic datasets for m6A target genes that are not disrupted at the RNA level but are decreased upon METTL3 depletion, and therefore promoted by m6A. We find that cytoskeleton reorganization and invasion genes that are associated with metastasis, are promoted in doxorubicin-treated cells compared to untreated (Table S1), and are m6A targets (meRIP) that are downregulated upon METTL3 depletion at the protein level but not RNA level (Fig. 4Gi, Table S3-S5). This is consistent with the fact that therapies like doxorubicin can induce epithelial to mesenchymal transition (EMT) and invasion of cancer cells (Kalluri and Weinberg, 2009; Mohammed et al., 2021). Therefore, we tested whether METTL3 was required for invasion of doxorubicin surviving cells. Consistently, we find that METTL3 depletion decreases invasion genes (Fig. 4Gi), such as Lyn and Eno2 (Choi et al., 2010), and consequently, decreases invasiveness of these cancer cells (Fig. 4Gii). Thus, Mettl3 increase upon therapy treatment, not only reduces proliferation via inhibition of cell cycle genes, but also increases invasion related genes, as a stress response via ISR in therapy-treated cells, enabling tumor persistence.

### Precluding non-canonical translation of METTL3 with ISRIB, enhances immune cell migration

Genes that are upregulated in doxorubicin-treatment and are promoted by m6A, include the DNA repair gene PARP, and the RNA Cytosine to Uridine editing enzyme, ADAR1 (Table S3-S5, Fig. S4B). PARP increases in doxorubicin-treated cells and in eIF2α phosphatase inhibitor, Sal003-treated cells (Fig. S4C), consistent with METTL3 increase. PARP is known to promote tumor survival and inhibit anti-tumor immunity (Pantelidou et al., 2019; Wanderley et al., 2022). Additionally, ADAR1 is enhanced (Fig. S4B-C), which can modify RNAs to prevent triggering anti-viral immune receptors (Chung et al., 2018; Ding et al., 2020; Kung et al., 2021; Lamers et al., 2019; Ramírez-Moya et al., 2020; Zipeto et al., 2016) and innate immune cell modulators that are also decreased by METTL3 (Fig 4Aii, Table S3-S5), leading to reduced anti-viral immune and cell death response in doxorubicin-treated cells. Together, this would disable anti-tumor immunity and can contribute to survival of such tumor cells; conversely, inhibition of METTL3 would activate immune response. First, we tested whether immune cell migration would be affected by METTL3 expression in doxorubicin-treated cells. Trans-well assays reveal that doxorubicin-treated MCF7 cells decrease immune cell migration of CD14+ monocytes from the top chamber to the bottom chamber containing such MCF7 cells; however, this was reversed and showed enhanced immune cell migration, if the MCF7 cells were treated with doxorubicin *and* ISRIB, or if METTL3 depleted cells were tested (Fig. 4H, S4D). These data suggest that m6A/METTL3 increase in chemo-treated cells disables immune cell migration, which can enable survival of such tumor cells.

### Therapy-treatment and METTL3 alter metabolic genes that can feedback regulate METTL3 levels

We find that several one-carbon metabolism genes that are needed to produce SAM (Cuthbertson et al., 2021; Green et al., 2019; Yang and Vousden, 2016; Zhu and Leung, 2020), the substrate for methylation, are m6A modified and regulated upon METTL3 depletion (Fig. 5Ai-ii, Table S3-S4). These genes, such as SHMT, are initially increased with doxorubicin treatment, and decreased at later times (24-48 h) of doxorubicin treatment (Fig. 5B until 18 h), co-incident with METTL3 increase. Consistent with the need for SAM for m6A modification of METTL3 mRNA for its translation upregulation (Fig. 3F-I), we find that addition of SHIN1, the inhibitor of SHMT (Cuthbertson et al., 2021; Guiducci et al., 2019; Herbig et al., 2002; Parsa et al., 2020; Pikman et al., 2022), reduces METTL3 levels (Fig. 5C). These data suggest that METTL3 regulates methylation via metabolism gene expression, and is regulated in turn by the metabolic state of the cells.

**Fig. 5.**
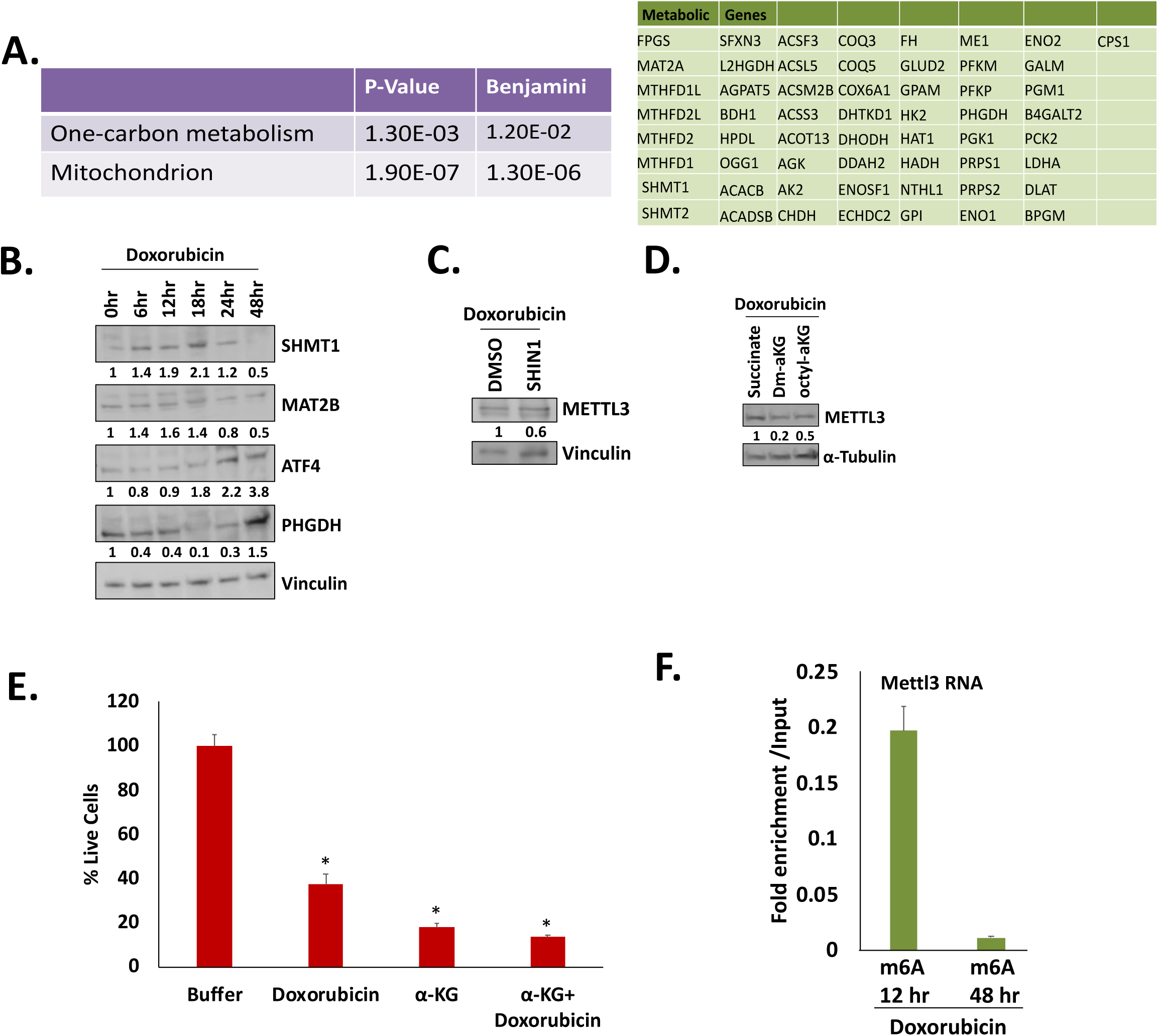
Metabolic genes that control methylation, are altered over time post-therapy, coincident with reduced m6A on METTL3 mRNA over time of doxorubicin treatment. **A.** Metabolism genes and one-carbon gene RNAs that are m6A modified (from Table S3-S4), and are regulated in doxorubicin-treated cells compared to untreated (from Table S1), and conversely in shMETTL3 cells compared to control shRNA MCF7 cells (from Table S3-S4). **B.** Western blot of metabolism genes over time of doxorubicin treatment: one-carbon metabolism genes (SHMT, MAT2B), ATF4 that increases PHGDH that can enhance α-ketoglutarate (αKG) that can lead to demethylation. Western blot analysis of impact on METTL3 levels upon addition of: **C.** SHIN1, the inhibitor of SHMT compared to buffer control, **D.** αKG compared to succinate or buffer control, and **E.** αKG on chemosurvival. **F.** m6A antibody immunoprecipitation at an early time-point after doxorubicin treatment (12 h where METTL3 increases, Fig. 1A) and at a later time (48 h where METTL3 levels do not increase, Fig. S1F), compared to IgG control, followed by qPCR analysis for METTL3 mRNA. See also Fig. S5, Table S3-5.

ISR-induced phospho-eIF2α translates ATF4 (Pakos-Zebrucka et al., 2016) that in turn, activates the PHDGH-mediated serine synthesis pathway that produces almost half of the cell’s α-ketoglutarate (αKG) (Li et al., 2021; Pacold et al., 2016). Consistent with early induction of phospho-eIF2α by doxorubicin treatment (Fig. 1A), we find ATF4 and PHDGH increase at later times of doxorubicin treatment (24-48 h doxorubicin treatment), after initial increase of phospho-eIF2α (Fig. 5B). αKG activates m6A demethylases (Kim and Lee, 2021; Su et al., 2018). SHMT is also blocked by PHGDH (Li et al., 2021; Pacold et al., 2016), and this is consistent with the increase of PHGDH and decrease of SHMT at later time points, decreasing SAM for methylation, revealing a potential feedback loop. This could reduce METTL3 mRNA methylation and reduce METTL3 translation as feedback regulation at later times (Fig. S1F). Consistently, addition of αKG but not succinate, reduces METTL3 levels (Fig. 5D), and decreases chemosurvival (Fig. 5E).

In accord with this temporal regulation of SAM and αKG enzymes, m6A antibody co-immunoprecipitates METTL3 RNA at early time points of 12 h when ATF4 and PHGDH levels are low, but one-carbon genes are high and not decreased yet (Fig. 5B) and METTL3 levels increase with this m6A modification of its mRNA (Fig. 1A, 3); however, the association of METTL3 mRNA with m6A antibody is reduced at 48 h when ATF4 and PHGDH are elevated (Fig. 5F) and METTL3 levels consistently decrease (Fig. S1F). These data suggest that therapy stress and m6A regulate enzymes that alter metabolites to control m6A on METTL3 mRNA; and thus, METTL3 levels, as feedback regulation.

## Discussion

Our previous studies revealed that chemoresistant cells, alter post-transcriptional gene expression to express as well as suppress specific genes, to enable chemosurvival (Lee et al., 2020). We find that therapy stress activates ISR and reduces mTOR activity; together, these two pathways inhibit both rate-limiting steps of canonical translation initiation. Our data reveal that the RNA m6A methyltransferase, METTL3, increases transiently in cancer cell lines and patient samples treated with chemotherapy (Fig. 1A, E-F, S1A-F, Table S1). This is observed with different chemotherapies in multiple cancer cell lines, in patient samples, and in pdx. Consistent with the enhanced METTL3, m6A on RNA is elevated in chemo-treated cells, but not in cells with METTL3 depletion (Fig. 1B, S1E). m6A modification of RNAs is known to alter gene expression (Jaffrey and Kharas, 2017; Ke et al., 2017; Liu et al., 2014; Meyer and Jaffrey, 2014; Roundtree et al.; Svitkin et al., 2017). Depletion of METTL3, or expression of a catalytically defective METTL3 compared to wildtype METTL3, decreases chemosurvival, indicating that the increased METTL3 and m6A modification on RNAs in chemo-treated cells, is needed for chemosurvival (Fig. 1D). These data suggest that therapy-treated cells inhibit canonical translation, and increase METTL3 as a stress adaptation, to regulate gene expression that contributes to chemosurvival.

Therapy-induced stress signaling leads to activation of ISR via eIF2α kinases such as PKR (Gandin et al., 2016; Harvey et al., 2019; Jewer et al., 2020; Lee et al., 2020), along with increased METTL3 (Fig. 1–2, S1–2). Consistently, METTL3 induction in chemotherapy-treated cells, can be mimicked by ISR inducers (Fig. 2A, S2A), such as poly I:C that activates PKR (Berlanga et al., 2006; Costa-Mattioli and Walter, 2020; Harding et al., 2003; Zyryanova et al., 2018). Furthermore, a phosphatase inhibitor, Sal003 that blocks dephosphorylation of eIF2α, also increases METTL3, indicating that eIF2α phosphorylation is needed (Fig. 2C). Consistently, inhibitors that override the effects of eIF2α phosphorylation, prevent METTL3 enhancement (Fig. 2D-E). These include Trazodone that is thought to affect the ternary complex (Harvey et al., 2019), or ISRIB that restores canonical ternary complex function, despite eIF2α phosphorylation (Costa-Mattioli and Walter, 2020; Sidrauski et al., 2013; Zyryanova et al., 2018). Consistent with elevated METTL3, m6A was amplified on RNA in cells with poly I:C, or Sal003, as in chemosurviving cells, along with augmented phospho-eIF2α, but not in cells treated with inhibitors that override ISR (Fig. 1B-C, 2B, S1G, S2G). Other signaling pathways like ATM that affects METTL3 (Zhang et al., 2020), may also contribute to regulation of METTL3. With eIF2α phosphorylation (Fig, 1-2, S1-2) and blocked mTOR activity (Fig. S1A, S2B-E) in doxorubicin-treated cells, canonical translation is reduced to allow recessive non-canonical mechanisms to translate specific mRNAs (Holcik, 2015; Spriggs et al., 2008; Walters and Thompson, 2016). Accordingly, METTL3 elevation is not observed at the RNA levels (Fig. 2F), suggesting translational control. This was confirmed by polysome analysis and nascent translation analysis that show increased polysome association and translation of METTL3, while other genes that are not needed in chemo-treated, arrested cells like KI67, are reduced on polysomes (Fig. 2G-I). These data indicate that chemotherapy induces the ISR pathway that causes phosphorylation of eIF2α, to enhance METTL3 via non-canonical translation in these conditions of reduced canonical translation.

Specific mRNAs can be recruited for non-canonical translation by distinct RNA binding protein complexes that connect with the translation machinery. We identified the RNA binding translation factor, eIF3 complex proteins (eIF3d, e, g, a, m, Fig. 3A, Fig. S3A-B, Table S2), as associated with METTL3 mRNA, by in vivo crosslinking-coupled biotin-tagged antisense purification of METTL3 mRNA. Consistently, depletion of eIF3a reduced METTL3 levels and chemosurvival, and overexpression of eIF3d enhanced METTL3 levels and chemosurvival (Fig. 3B-E). EIF3 can promote canonical translation via recruitment of the ribosome to mRNAs, as well as non-canonical translation via direct association with m6A-modified mRNAs and other specific mRNAs (Barbieri et al., 2017; Edupuganti et al., 2017; Meyer, 2019; Meyer et al., 2015b). Therefore, we tested METTL3 5′UTR reporters, and identified that the 5′UTR was sufficient to confer translation increase upon doxorubicin treatment but not when the GAC motif upstream of the ATG was mutated (Fig. 3F-H). METTL3 is known to have such 5′UTR m6A modification GAC sites that respond to non-canonical translation under heat shock stress and Torin 1 inhibition (Zhou et al., 2018). Furthermore, we find that reducing m6A via METTL3 depletion, also reduces METTL3 5′UTR Luciferase reporter translation (Fig. 3I). METTL14 mRNA that is co-regulated with METTL3, also binds eIF3, and remains to be explored. These data suggest that therapy-induced eIF2α phosphorylation, reduces canonical translation, to enable specialized translation on METTL3 mRNA via its m6A-modified 5′UTR that binds eIF3.

m6A is known to mediate post-transcriptional regulation, including RNA decay of its targets (Jaffrey and Kharas, 2017; Ke et al., 2017; Liu et al., 2014; Meyer and Jaffrey, 2014; Roundtree et al.; Svitkin et al., 2017). Accordingly, we find in doxorubicin-treated cells, m6A predominantly decreases cell cycle-related genes at the RNA level (Fig. 4Ai-ii, Table S3 for m6A-modified RNAs and RNAs regulated by METTL3 depletion, Table S4 for protein changes upon METTL3 depletion, Table S5 for common genes from Table S3-S4). This is consistent with enabling chemosurvival, where the cell cycle must be inhibited to avoid cell death due to chemotherapy that targets proliferation. Accordingly, we find that METTL3 deletion decreases chemosurvival (Fig. 1D), with concomitant increase in cell cycle genes (Fig 4Aiii). Thus, chemo-treated cells may survive therapy, via ISR stress signaling, triggered by chemotherapy, to increase METTL3 to downregulate proliferation gene expression to suppress chemosensitivity.

Apart from cell cycle genes, METTL3 downregulates anti-viral immune response genes (Fig. 4Aii, B-C, Table S3-S5), including PKR and RIG-I/DDX58, the viral RNA/pattern recognition receptor that mediates type-1 interferon response when unmodified non-self RNA (that appears non-self) is present (Boelens et al., 2014; Loo and Gale, 2011) (Dempoya et al., 2012; Majzoub et al., 2019; Streicher and Jouvenet, 2019). m6A and other modifications are known to mark RNAs as self-RNAs to prevent triggering anti-viral immune response via RIG-I (Chen et al., 2019; Durbin et al., 2016; Karikó et al., 2005) that can lead to inflammation-mediated tumor cell death and innate immune response to attract immune cells (Dempoya et al., 2012; Loo and Gale, 2011). Consistently, METTL3 depleted cells increase anti-viral response and downstream STAT1 and CASP9 signaling (Fig. 4D); conversely, poly I:C triggers a reduced anti-viral response in METTL3 overexpression cells compared to a catalytic mutant, with decreased expression of PKR and RIG-I due to increased m6A (Fig. 4E). The increase in METTL3 and thus m6A on RNAs, may be the mechanism to prevent this induced cell death, both by methylating endogenous RNAs that then do not trigger an anti-viral immune response as they are recognized as self-RNAs, as well as by downregulating anti-viral immune response genes and cell cycle genes.

METTL3 also promotes translation of specific gene categories that are modified by m6A (Table S3-S5). METTL3 depletion decreases invasion genes; consistently, METTL3 depleted cells show reduced invasion (Fig. 4Gi-ii). This is consistent with the fact that therapies like doxorubicin can induce EMT and invasion (Choi et al., 2010; Kalluri and Weinberg, 2009; Mohammed et al., 2021). Additionally, METTL3 depletion reduces the expression of the DNA repair gene, PARP, as well as the RNA editor, ADAR (Fig. S4B-C); these increase in chemo-treated cells and would increase chemosurvival and immune evasion, as recently observed with m6A regulation of ADAR1 (Terajima et al., 2021). These findings along with downregulation of anti-viral immune genes, suggested that immune cell migration would be reduced, to avoid immune response-induced cell death. In accord, a trans-well assay shows decreased monocyte migration toward doxorubicin-treated MCF7 cells in the bottom chamber, compared to untreated cells; in contrast, monocyte migration is enhanced when METTL3 is depleted or when MCF7 chemo-treated cells were co-treated with ISRIB to reduce METTL3 (Fig. 4H, S4D). These data support that METTL3 is elevated in chemo-treated cells to preclude innate immune response.

Our data suggest that chemotherapy induces ISR via eIF2α phosphorylation, and reduces conventional translation, to enable eIF3-mediated, non-canonical translation upregulation of METTL3. We find that m6A modification on mRNAs by therapy-induced METTL3, leads to post-transcriptional downregulation of genes that need to be suppressed to escape chemotherapy, as well as promotes specific genes that enable survival (Fig. 4, S4). This suggests that non-canonical translation of METTL3 is a potential vulnerability of chemoresistant cells, which can be targeted to limit tumor survival. Consistently, we find that overriding the effect of eIF2α phosphorylation, with trazodone (Harvey et al., 2019) or ISRIB (Costa-Mattioli and Walter, 2020; Sidrauski et al., 2013), prevents METTL3 increase (Fig. 2D-E); and accordingly, dysregulates target genes and reduces chemosurvival (Fig. 4E-F). Depletion of METTL3 reduces chemosurvival but addition of trazodone did not reduce chemosurvival further, indicating that METTL3 and eIF2α regulation are in the same pathway that causes chemoresistance (Fig. S4A). These data suggest potential inhibitors that can be combined to improve the efficacy of chemotherapy, by reducing non-canonical translation of METTL3.

The effects of doxorubicin or eIF2α phosphorylation are transient (6-18 h, Fig. S1F) and METTL3 levels are subsequently restored, suggesting that METTL3 increases as a transient stress response. METTL3 uses the donor for methylation, SAM, to increase its own production, and to regulate target genes for chemosurvival. METTL3 alters one-carbon metabolism genes (Fig. 5A, Table S3-S4) that produce SAM, which would reduce methylation over time and thereby could feedback regulate METTL3 translation. Therapy-induced ISR mediates eIF2α phosphorylation that increases METTL3 via non-canonical translation by its m6A 5′UTR site. Therapy also induces regulators that promote SAM production (Leclerc and Rozen, 2008; Zhu and Leung, 2020). We find increased SHMT, and MAT2B (Fig. 5B) that produce SAM (Cuthbertson et al., 2021; Green et al., 2019; Yang and Vousden, 2016; Zhu and Leung, 2020), at early time points till 18 h where METTL3 also increases (Fig. 5B). Consistently, we find that co-treatment with doxorubicin and SHIN1, an inhibitor of SHMT, leads to decreased METTL3 (Fig. 5C). ATF4 induced over time by eIF2α phosphorylation, can promote PHGDH, which enhances αKG (Li et al., 2021; Pacold et al., 2016) that activates m6A demethylases (Kim and Lee, 2021; Su et al., 2018); this would then downregulate m6A on METTL3 mRNA and thus METTL3 translation at later times of therapy treatment. In accord, we find METTL3 decreases while PHGDH, increases at 24-48 h where ATF4 accumulates (Fig. S1D, 5B). Consistently, we find that co-treatment with doxorubicin and αKG reduces METTL3 levels and chemosurvival (Fig. 5D-E). Additionally, increased METTL3 downregulated not only cell cycle genes for chemosurvival but also PKR (Fig. 4B-D) that can curb eIF2α phosphorylation over time, which would reduce METTL3 translation. Thus, RNA metabolism by m6A is regulated by stress signals triggered by doxorubicin treatment, which in turn alters metabolite levels by affecting enzymes that regulate SAM, αKG, and eIF2α phosphorylation, leading to temporal m6A regulation to avoid anti-tumor activity by cell cycle and anti-viral genes. Supporting this model of temporal regulation of METTL3 by m6A over time of therapy treatment (Fig. S5), we find that METTL3 mRNA associates significantly with m6A antibody at early (12 h) but not at later times (48 h) of doxorubicin treatment (Fig. 5F). Together, our data reveal that therapy induces non-canonical translation of METTL3 that leads to m6A regulation for tumor survival—which can be blocked, by interfering with therapy-induced non-canonical translation or metabolic change, to curtail tumor survival.

## Materials and Methods

### Cell Culture

THP1 cells were cultured in RPMI1460 media supplemented with 10% fetal bovine serum (FBS), 2 mM L-Glutamine, 100 µg/mL streptomycin and 100 U/ml penicillin at 37°C in 5% CO_2_. MCF7, BT549 cells were grown in DMEM supplemented with 10% fetal bovine serum (FBS), 2 mM L-Glutamine, 100 µg/mL streptomycin and 100 U/ml penicillin at 37°C in 5% CO_2_. Cells were treated with indicated concentrations of therapies for indicated periods of time. All cell lines and monocytes (CRL9855) were obtained from ATCC (Sykes et al., 2016). Cell lines were tested for Mycoplasma (Promega) and authenticated by the ATCC Cell Authentication Testing Service (Lee et al., 2020).

### Tumorspheres, patient samples, and animal samples

Hormone positive and triple negative patient samples were obtained were from MGH in accord with IRB approved protocols (AL, LE, SBK, SV).

Hormone positive patient sample cells were isolated and relabeled as BT 29 and BT30, used in figures 1E, 2C-E, S2E. Tumorspheres were grown in Matrigel (#CB40230C, Corning) as described (Lee et al., 2007; Muranen et al., 2012; Weigelt et al., 2014). Briefly, Human breast tissues were dissociated into single-cell suspension by enzymatic digestion using Milteny Biotec Tumor dissociation Kit human (cat#130-095-929). The enzyme mix was prepared as per the manufacturer’s guidelines. Tissues were washed with sterile cold PBS and then fat, fibrous, and necrotic tissues were removed. Tumor tissue (0.2-1.0 g) was chopped into small pieces (2-4mm) using sterile surgical blades and transferred into the gentleMACS C tubes containing the enzyme mix (enzyme H 200ul + enzyme R 100ul + enzyme A 25ul). Tube C was closed tightly and attached upside down onto the sleeve of gentleMACS™ Octo Dissociator with Heaters. An appropriate gentleMACS™ Program (37C_h_TDK_3 for tough tissues) was run and after the termination of the program, tubes were detached from the gentle MACS Dissociator and briefly centrifuged to collect samples at the bottom of the tube. Samples were resuspended and the cell suspension was passed through the MACS SmartStrainer (30 µm or 70 µm) placed on a 50 mL tube. MACS SmartStrainer (30 µm or 70 µm) was washed with 20 mL of DMEM media and the tubes were centrifuged at 300xg for 7 min and the supernatant was aspirated completely. Cell pellets were resuspended in DMEM and about 3000 cells were transferred into one tube and the rest in another tube and centrifuged at 300xg for 7 min. About 2000-3000 cells were resuspended into 200ul - 400ul of Matrigel that was thawed on ice for 30min and plated as 20ul drops on two pre-warmed 10cm plates without any bubbles and then plates were carefully placed at a 37degree incubator for 20 min to let the Matrigel polymerize and then media was added. Every two days media was changed, and tumorspheres were allowed to grow for 7-10 days then one plate was treated with doxorubicin for 24 hours before tumorspheres were harvested for western blot. The other set of cells was plated in a 10 cm plate and allowed to grow for propagation.

PDX samples were obtained from SBK and LE. Animals were housed and treated in accordance with protocols approved by the Subcommittee on Research Animal Care at the MGH. Immunodeficient NOD scid gamma mice were implanted with patient-derived breast tumors (Koh et al., 2021). Once tumors reached a volume of ~100 cubic millimeter, mice were treated with vehicle (saline) or 2 mg/kg doxorubicin (Selleck Chemicals) for 1 h. Tumors were excised and snap-frozen before being processed for subsequent protein-based analyses.

### Plasmids

GIPZ plasmids expressing shRNAs against human METTL3 (from MGH MPL (pLKO.1 shRNA-Mettl3(B12) GCCAAGGAACAATCCATTGT), and control vector expressing miR30a primiR sequences (RHS4750), shRNA against eIF3A (V3THS_386738 ATTGTAGTACATTAAATCT) were obtained from Open Biosystems-Dharmacon. Stable cell lines were constructed as described previously (Bukhari et al., 2016; Le et al., 2016). Renilla was obtained and used previously (Vasudevan and Steitz, 2007). The 5′UTR of METTL3 was amplified (with primers flanked by NheI restriction sites). CX plasmid (Bukhari et al., 2016) and 5′UTR containing amplicon was digested with NheI (NEB) followed by ligation with T4DNA ligase (NEB). The 3′UTR of METTL3 was amplified using primers with Not I sites and cloned into CX plasmid as described earlier (Bukhari et al., 2016). GAC mutation in the 5′UTR was constructed using Quikchange (Agilient-Stratagene). Ligation mixes were transformed in *E. Coli* cells. Plasmids were purified from *E. Coli* cells post transformation. Positive clones were confirmed by sequencing.

### Polysome profiling and microarray

30 × 10^6 cells were grown for each sample, treated with 100ug/ml of cycloheximide for 5 min, prior to harvesting cells on ice. Sucrose was dissolved in lysis buffer containing 100 mM KCl, 5 mM MgCl_2_, 2 mM DTT and 10 mM Tris-HCl (pH 7.4). Sucrose gradients from 10% to 55% were prepared in ultracentrifuge tubes (Beckman) as previously described(Truitt et al., 2015; Vasudevan and Steitz, 2007). 100 ug/ml of cycloheximide was added to lysis buffer and sucrose solutions in case of cycloheximide treatment of cells prior to harvesting. Harvested cells were rinsed with ice-cold PBS and resuspended in lysis buffer with 1% Triton X-100 and 40 U/mL murine (New England Biolabs) for 20 minutes with intermittent tapping on ice. After centrifugation of cell lysates at 12,000 × g for 1 h 20 minutes, supernatants were loaded onto sucrose gradients followed by ultracentrifugation (Beckman Coulter Optima L90) at 32,500 × rpm at 4 °C for 80 min in the SW40 rotor. Samples were separated by density gradient fractionation system (Biocomp Piston Gradient Fractionation). RNAs were purified from heavy polysome fractions and whole cell lysates. The synthesized cDNA probes from WT Expression Kit (Ambion) were hybridized to Gene Chip Human Transcriptome Array 2.0 (Affymetrix) and analyzed by the Partners Healthcare Center for Personalized Genetic Medicine Microarray and BUMC facilities. Gene ontology analysis for differentially expressed translatome or proteome was conducted by DAVID 6.7 tools (Huang et al., 2009). Molecular signatures enriched were identified by Gene Set Enrichment Analysis (GSEA) (Subramanian et al., 2005a).

### Western blot analysis

Cells were collected and resuspended in lysis buffer containing 40 mM Tris-HCl (pH 7.4), 6 mM MgCl_2_, 150 mM NaCl, 0.1% NP-40, 1 mM DTT, 17.5 mM ß-glycerophosphate, 5 mM NaF and protease inhibitors. Cell lysates were prepared by sonication and low speed centrifugation, after Micrococcal nuclease (NEB) treatment to remove RNA and DNA, and were heated at 95°C with 200 mM DTT and 1X SDS loading dye for 10 min. Samples were loaded onto 4%-20% gradient SDS-PAGE (Bio-Rad) or 16% SDS-PAGE (Invitrogen), transferred to PVDF membranes and processed for immunoblotting. Antibodies against METTL3 (15073-1-AP), m6A (MA5-33030), Actin (MAB1501), Tubulin (05-829), Vinculin (26520-1-AP), ATF4 (10835-1-AP), were from Protein Tech, Thermo-Fisher, and Millipore; PKR (3072S), P-PKR (2611S), EIF2α (9722S), P-EIF2α (3597S), were from Cell Signaling; ATF4 (ab23760) was from Abcam. All antibodies are listed in Table S6.

### Mass Spectrometry for proteome analysis

Multiplex quantitative proteomics analysis (tandem-mass-tag (TMT) spectrometry) was conducted, as previously (Bukhari et al., 2016), from cell lines created that were treated as described.

#### Cell pellets

Cell pellets were lysed (75mM NaCl, 50mM HEPES pH 8.5, 10mM Sodium pyrophosphate, 10mM Sodium Fluoride, 10mM B-Glycerophosphate, 10mM Sodium Orthovanadate, Roche complete mini EDTA free protease inhibitors, 3% SDS, 10mM PMSF), reduced and alkylated and underwent tryptic digest as previously described (Edwards and Haas, 2016). 50 µg of the resulting peptides were subsequently labeled using TMT-11plex reagents (Thermo-Scientific) according to manufacturer’s instructions. Labeled samples got combined and fractionated using a basic reversed phase HPLC (Edwards and Haas, 2016).

#### Immunoprecipitates

Eluate was reduced and alkylated and underwent tryptic digest as previously described (Edwards and Haas, 2016; Perugino et al., 2019). 10 ug of these peptides underwent subsequent labeling with TMT-11plex reagents (Thermo-Scientific) according to manufacturer’s instructions. Combined labels were desalted.

#### Mass spectrometry and Bioinformatics

The resulting fractions from the cell pellets and the desalted TMT sets from the IPs were analyzed in an 3h reversed phase LC-MS2/MS3 run on an Orbitrap fusion Lumos or an Orbitrap Fusion (Thermo-Scientific). MS3 isolation for quantification used Simultaneous Precursor Selection (SPS) as previously described (Erickson et al., 2015; McAlister et al., 2014; Ting et al., 2011). Proteins were identified based on MS2 spectra using the Sequest algorithm searching against a human data base (Uniprot 2014) (Eng et al., 1994) using an in house built platform (Huttlin et al., 2010). Search strategy included a target-decoy database-based search in order to filter against a false-discovery rate (FDR) of protein identifications of less than 1% (Elias and Gygi, 2007).

### Quantitative RT-PCR

Total RNA was extracted using proteinase K buffer and TRIzol (Invitrogen) as performed previously (Bukhari et al., 2016). The cDNA was synthesized from 1 μg of RNA using M-MuLV Reverse Transcriptase (NEB) and random hexamer primer (Promega). qPCRs were run on LightCycler® 480 Instrument II (Roche) using 2 X SYBR green mix (Bio-rad). All primers used are listed in Table S6.

### RNA modification analysis

Total RNA was isolated using Trizol as per manufacturer’s instructions. Isolated RNAs were cleaned using RNeasy kit (Qiagen). Poly(A) containing RNAs were separated from the RNA pool using a Poly(A) mRNA isolation system IV (PolyATract, Promega), followed by Ribozero purification, and a microspin column to remove RNAs <200nt to eliminate tRNAs. The RNAs were tested by dot blot analysis and 2D-TLC as performed previously (Chen et al., 2020).

For dot blot analysis, the RNA is loaded on a positive-charged Nylon66 membrane (Biodyne B transfer membrane, 0.45 μm, 60209), and cross-linked by a UV Stratalinker 2400 (Stratagene; La Jolla, CA, USA) at 1200 μJ twice. Then, the membrane is washed in TBST for 10 min. Primary m6A antibody was diluted 1:100 in 5% non-fat milk in TBST. After overnight incubation at 4L°C, the membrane was washed three times in TBST for 10 min. Secondary antibody was diluted 1:2000 in 5% non-fat milk in TBST and then washed three times.

2D-TLC was performed as described previously(Grosjean et al., 2004; Zhong et al., 2008). Approximately 500 ng of RNA was digested with RNase If and RNase T2 (Cat#: M0243L, New England Biolabs/NEB and Cat#: LS01502, Worthington Biochemical Corp.), followed by labeling with γ-P^32^-ATP and T4 polynucleotide kinase (Cat#: M0201L, NEB). The reactions were further processed and digested with P1 Nuclease (Cat#: M0660S, NEB). In vitro-transcribed 4B mRNA (firefly luciferase reporter (Bukhari et al., 2016)), with or without m6A,was run in parallel as controls. The reactions were loaded on cellulose TLC plates (20L×L20Lcm, EMD Millipore™ Precoated TLC and PLC Glass Plates, Cat#: M1057160001, ThermoFisher Scientific) and developed in two solvent systems: solvent A with isobutyric acid: 0.5LM NH_4_OH (5:3Lv/v; Cat#: AAL04038AP, AC423305000) for the first dimension and solvent B with phosphate buffer/ammonium sulfate/*n*-propanol (100/60/2 (v/w/v)) for the second perpendicular dimension and then exposed to film. The spots were quantified with ImageJ software and normalized for total input cpms for comparison between samples.

### Nascent translation level analysis

Global translation was measured by metabolic labeling for a short period followed by PAGE and scintillation analysis (Vasudevan and Steitz, 2007). MCF7 cells with or without doxorubicin treatment, were grown in normal DMEM medium to prevent additional cellular stress from methionine-free medium. Nascent translation analysis using L-azidohomoalanine (AHA), an amino acid analog of methionine containing an alkyne moiety that can be biotinylated by click-it chemistry (ThermoFisher), for nascent protein translation labeling followed by SDS-PAGE and HRP-streptavidin Western analysis; AHA was added as described in the manufacturer’s protocol. After incubation at 37 °C for 45 min, cells were washed once with PBS and lysed in buffer (40 mM Tris-HCl (pH 7.4), 6 mM MgCl_2_, 150 mM NaCl, 0.1% NP-40, 1 mM DTT and protease inhibitors). The lysate was first separated by electrophoresis on an SDS-PAGE gel, then transferred to a nitrocellulose membrane by Western Blotting with anti-biotin antibody. These were quantified by ImageJ.

### In vivo crosslinking and Immunoprecipitation

In vivo crosslinking using 0.3% formaldehyde and UV 254 nm (using a Stratalinker), and nuclear-cytoplasmic separation were done as described earlier (Bukhari et al., 2016; Lee et al., 2020). Briefly, around 10-15 × 10^6 cells were harvested on ice followed by washing with cold PBS. Cell pellets were resuspended in hypotonic buffer (10 mM Tris (pH=8), 1.5 mM MgCl_2_, 10 mM KCl). Cells were lysed using a syringe with 25G × 5/8 precision glide needle and a syringe were used to lyse the cell by repetitive passes through the needle (up to 5-10 times to ensure lysis) or, alternatively, a douncer was used to appropriately lyse the cells. The decanted supernatant (cytoplasmic fraction) was centrifuged at 2000 rpm for 10 min. at 4**°**C. The supernatant was again decanted at 2000 rpm for 10 mins producing the final cytoplasmic fraction. The remaining pellet was washed with 10 times the pellet volume with Buffer B (20 mM Tris-HCl pH 8.0, 25% Glycerol, 1.5 mM MgCL_2_, 0.2 mM EDTA pH 8.0, 20 mM KCl). After removal of the wash buffer, the pellet was again suspended in 10 times the pellet volume of Buffer B and 5 times the pellet volume of Buffer C (20 mM Tris-HCl pH 8.0, 25% Glycerol, 1.5 mM MgCL_2_, 0.2 mM EDTA pH 8.0, 1.2 M KCl) was added and the solution was mixed by vortex before a 45 min incubation with nutation at 4**°**C. If the cells had been crosslinked, the nuclear fraction was sonicated for 5s then 30s cooling on ice (6 times, 90% duty cycle, output control at 2) and DNase I treated before being centrifuged at 2000 rpm for 10 min at 4**°**C. The decanted supernatant was collected as the nuclear fraction. The supernatant was mixed with the cytoplasmic fraction. Where mentioned, cytoplasmic fraction was used. The lysates were pre-cleared of non-specific binders by incubating with Protein G (Santa Cruz Biotechnology) beads and IgG. The pre-cleared lysates were incubated with antibodies and IgG (control) overnight in buffer (40 mM HEPES, 100 mM NaCl, 6 mM MgCl2, 0.025% NP-40, 1 mM DTT, 10% glycerol, 1mM PMSF). Lysates were then incubated with equilibrated and blocked Protein G beads for 2 h. Beads were then pelleted and washed 4 times with RIPA buffer. Beads were then either used to analyze proteins or RNA after using heat to break Schiff’s linkages from formaldehyde, followed by proteinase K digestion buffer treatment for the fractions for RNA analysis, and RNase treatment for the fraction for protein analysis as described earlier (Bukhari et al., 2016; Lee et al., 2020).

### DEAE (diethylaminoethanol) fractionation and RNA antisense purification

DEAE fractionation was performed with in vivo formaldehyde crosslinked extracts for further purification with RNA antisense against METTL3 as developed previously (Vasudevan and Steitz, 2007). Cell lysates were incubated with equilibrated DEAE beads for 2 hr in buffer (40 mM HEPES, 6 mM MgCl_2_, 2 mM DTT, 10% glycerol, 150 mM NaCl). After collection of the flowthrough, beads were incubated with wash buffer of increasing salt concentrations (40 mM HEPES, 6 mM MgCl_2_, 2 mM DTT, 10% glycerol, with NaCl ranging from 250 mM-500 mM-750 mM-1000 mM-2000 mM). RNA was isolated from each salt fraction. Amount of RNA in each fraction was analyzed by qRT-PCR, normalized to input levels to find those fractions containing METTL3 mRNA by qPCR. These fractions were pooled, DNase treated, and pre-cleared for 1 h over avidin beads to remove endogenous biotinylated proteins. This was then subjected to METTL3 or scrambled N biotinylated antisense oligo purification overnight. METTL3 shRNA sequence that showed efficient knockdown was used to design the METTL3 antisense as this showed that the site was available for base pairing in the cell, which was tagged with biotin (IDT). This was then enriched over streptavidin beads, washed 4 times with RIPA buffer, and the beads were then split into half which were either subjected to RNA elution by proteinase K digestion or to protein elution by RNase One treatment. The crosslinks in these eluates were disrupted by heat inactivation in proteinase K buffer as described previously (Vasudevan and Steitz, 2007).

### Luciferase Assay

Plasmids containing firefly luciferase reporters (Kearse et al., 2019) were co-transfected using Lonza Nucleofector (Bukhari et al., 2016) with Renilla luciferase or with Lipofectamine 2000 (Thermo) or MIRUSBIO reagents in MCF7, control and METTL3 shRNA stable cell lines. Transfected cells were then grown under conditions of doxorubicin or buffer treatment. Cells were harvested, washed, and lysed in 1X passive lysis buffer (Promega). Luciferase activity in the lysates was analyzed using Luciferase Assay System (Promega) as per manufacturer’s instructions and as conducted previously (Vasudevan and Steitz, 2007).

### Cell Migration Assay

Cell migration assay was performed as previously described (Le et al., 2016). Trans-well chambers (8 μm pore, Corning) were pre-equilibrated with serum-free media. CRL9855 monocytes (2 × 10^4^/chamber) that were pre-stained for 1 hr per manufacturer’s instructions with Far Red Cell Trace dye (Thermo Fischer Scientific), were placed in the top chamber, and 700 μl of MCF7 cells were placed in the bottom chamber. The chambers were incubated at 37°C for 18 hr in 5% CO_2_. Cells on the upper surface of the filter were removed with a cotton swab. Migrated stained monocyte cells were observed in the bottom chamber and visualized using a microscope. Microscope images were taken, and the numbers of migrated cells were determined.

### Inhibitors and chemicals

Doxorubicin, Gemcitabine, Paclitaxel, Cytarabine (AraC) (Kojima et al., 2005), Trazodone, and Trans-ISRIB (Costa-Mattioli and Walter, 2020; Zyryanova et al., 2021), were obtained from Cayman Chemicals and used as described. Sal003 10uM was obtained from selleckchem. Cells were treated with 1 µM trans-ISRIB or 5 µM Trazodone for 18-24 h. For cell viability that was measured by Trypan blue staining and cell counts (Lee et al., 2020), cells were treated individually or with a combination of chemotherapies and 1 µM trans-ISRIB or trazodone for 18-24 h.

### GO, GSEA analysis

Multiple Em for Motif Elicitation (MEME) software was used to search for 5′UTR elements as described earlier (Bailey and Elkan, 1994; Lee et al., 2020), and by looking for RRACH motifs, using 5’UTR sequences from Genbank. Gene ontology (GO) analysis for differentially expressed translatome or proteome was conducted by DAVID 6.7 tools (Huang et al., 2009) as described earlier (Lee et al., 2020) with our datasets. Gene Set Enrichment Analysis (GSEA) (Subramanian et al., 2005b) was performed as described earlier (Lee et al., 2020) with our datasets.

### Statistical analyses

Each experiment was repeated at least 3 times. No statistical method was used to pre-determine sample size. Sample sizes were estimated on the basis of availability and previous experiments (Bukhari et al., 2016). No samples were excluded from analyses. Statistical tests were conducted for each figure. SEM (standard error of mean) values are shown as error bars in all figures. P-values less than 0.05 were indicated with an asterisk. E-values were used for the statistical significance in the motif analysis. Statistical analyses for the datasets were performed as described. For the microarray data: signed linear fold changes were used. (e.g., +2 = 2-fold higher in doxorubicin treated than in untreated control; −2 = 2-fold lower in METTL3 KD than in control). Paired t tests (including replicate as a covariate) were performed for each gene between experimental groups to obtain a t statistic and p value for each gene. t statistics and nominal p values for each effect in the two-factor linear model, and t statistics and nominal p values for each pairwise comparison were generated. Benjamini-Hochberg False Discovery Rate (FDR) correction was applied across all genes to obtain FDR-corrected p values (FDR q values), which represent the probability that a given result is a false positive based on the distribution of all p values on the array. FDR q (filtered) values were recomputed separately for the two-factor model or each pairwise comparison, across only the genes that were expressed above the median value of at least as many samples as in the smallest experimental group involved in the model/comparison. NES is normalized enrichment score for GSEA. For mass spectrometry data in other tables: SD-standard deviation, SEM-standard error of mean, CI-confidence interval, p-value derived by student’s t-test using Excel.

## Author Contributions

SIAB lead and conducted the research; SST, CD, PC, KQW, JS, RM, EP, RE, SL, VB, and YL, contributed to the data; JK, RM (Morris), and WH conducted proteomics; SBK, LE, AL provided pdx and patient samples; SV lead and supervised the project, and wrote the manuscript. We thank Partners Healthcare Center for Personalized Genetic Medicine and BUMC facilities for microarray data.

## Data Availability

Raw datasets will be made available on the public repository, as supplementary tables or on GEO, and all materials will be made available publicly on publication and on request.

## Supplemental Figure legends

**Fig. S1.**
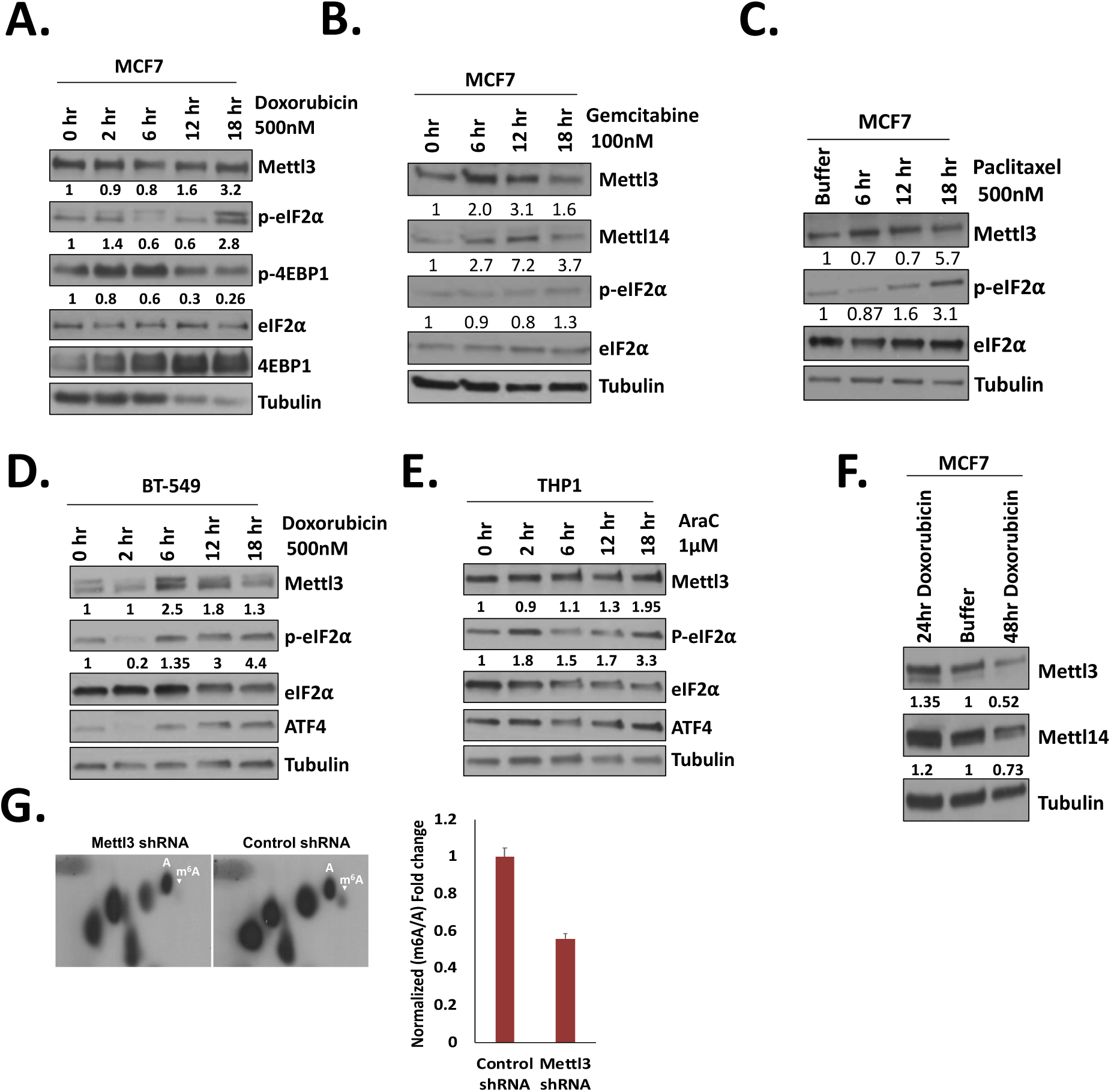
Western blots of MCF7 cells show increase of METTL3 over time, when **A.** treated with 500nM doxorubicin, **B.** treated with 100nM of gemcitabine, and **C.** treated with 500nM of paclitaxel. **D.** BT549 cells treated with 500nM doxorubicin, and **E.** THP1 cells treated with 1µM Cytarabine or AraC, over time show increased METTL3 and eIF2α phosphorylation. **F.** Western blot: METTL3 increase with 500nM doxorubicin treatment is a transient stress response and decreases after 24-48 h of treatment shown in MCF7 cells. **G.** 2D-TLC of rRNA and tRNA depleted, purified RNA from 500nM doxorubicin-treated control shRNA cells compared to METTL3 shRNA cells with graph of m6A quantified over total A. Data are average of 3 replicates +/−SEM. Tubulin serves as the loading control. See also Fig. 1, Table S1.

**Fig. S2.**
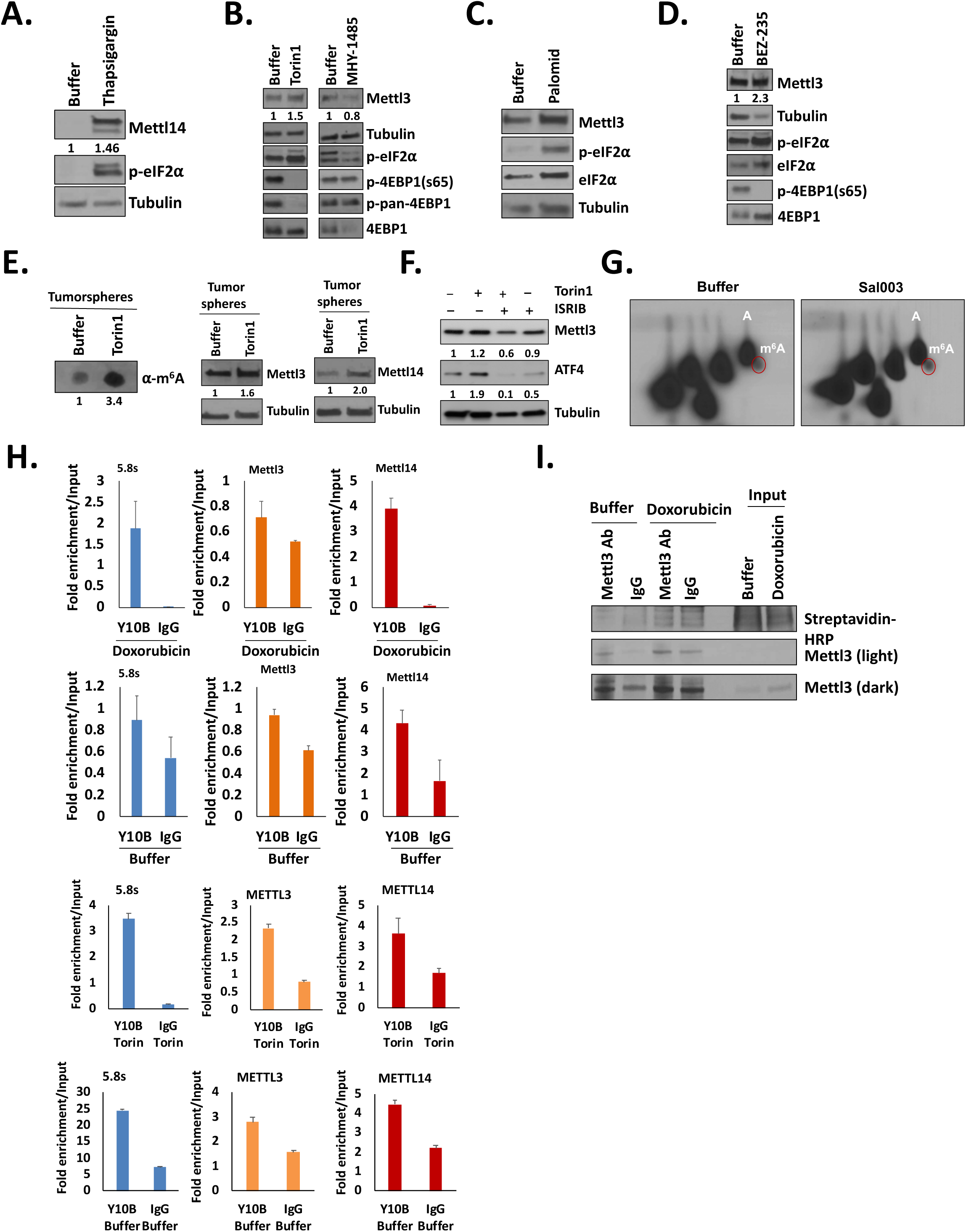
Western analyses of MCF7 cells treated with **A.** Thapsigargin (1uM for 1 day) to induce eIF2α phosphorylation, **B.** with 250nM Torin1 (2 days) to block mTOR and induce 4EBP dephosphorylation that coincides with eIF2α phosphorylation, to test METTL3 increase, as well as with mTOR activator, MHY1485 (2uM for 2 days), to inhibit mTOR and reduce 4EBP dephosphorylation and eIF2α phosphorylation, **C.** with 20uM Palomid (1 day) as another drug to block mTORC1 and mTORC2, and **D.** with 250nM BEZ235 (2 days) to dually block PI3K and mTOR, and induce 4EBP dephosphorylation and eIF2α phosphorylation. **E.** m6A dot blot analysis with 250nM Torin 1 (2 days) treatment of tumorspheres of tRNA (200nt cutoff microspin column purification) and ribosomal RNA depleted (ribo-zero purification) RNA from hormone-positive breast cancer patient sample BT30, where METTL3 and METTL14 increase as shown by Western blot analysis. **F.** 2D-TLC analysis of ribosomal RNA depleted, poly(A)-selected RNA from 10uM Sal003-treated MCF7 cells compared to buffer-treated cells, with nucleosides A and m6A marked. **G.** Y10B immunoprecipitation from 500nM doxorubicin (1day) treated cells to verify mRNAs with polysome association, followed by qPCR analysis of METTL3 and METTL14 mRNA and control 5.8S rRNA. Y10B immunoprecipitation from torin 1 (250nM for 2 days) treated MCF7 cells to verify mRNAs with polysome association, followed by qPCR analysis of METTL3 and METTL14 mRNA and control 5.8S rRNA. **H.** Nascent amino acid labeling and immunoprecipitation of METTL3, followed by Western analysis of nascently-translated and thus labeled METTL3 with Streptavidin-HRP and METTL3 antibodies to verify immunoprecipitation shown in Fig. 2I. Actin and Tubulin are loading controls for Western blots. Data are average of 3 replicates +/−SEM. See also Fig. 2.

**Fig. S3.**
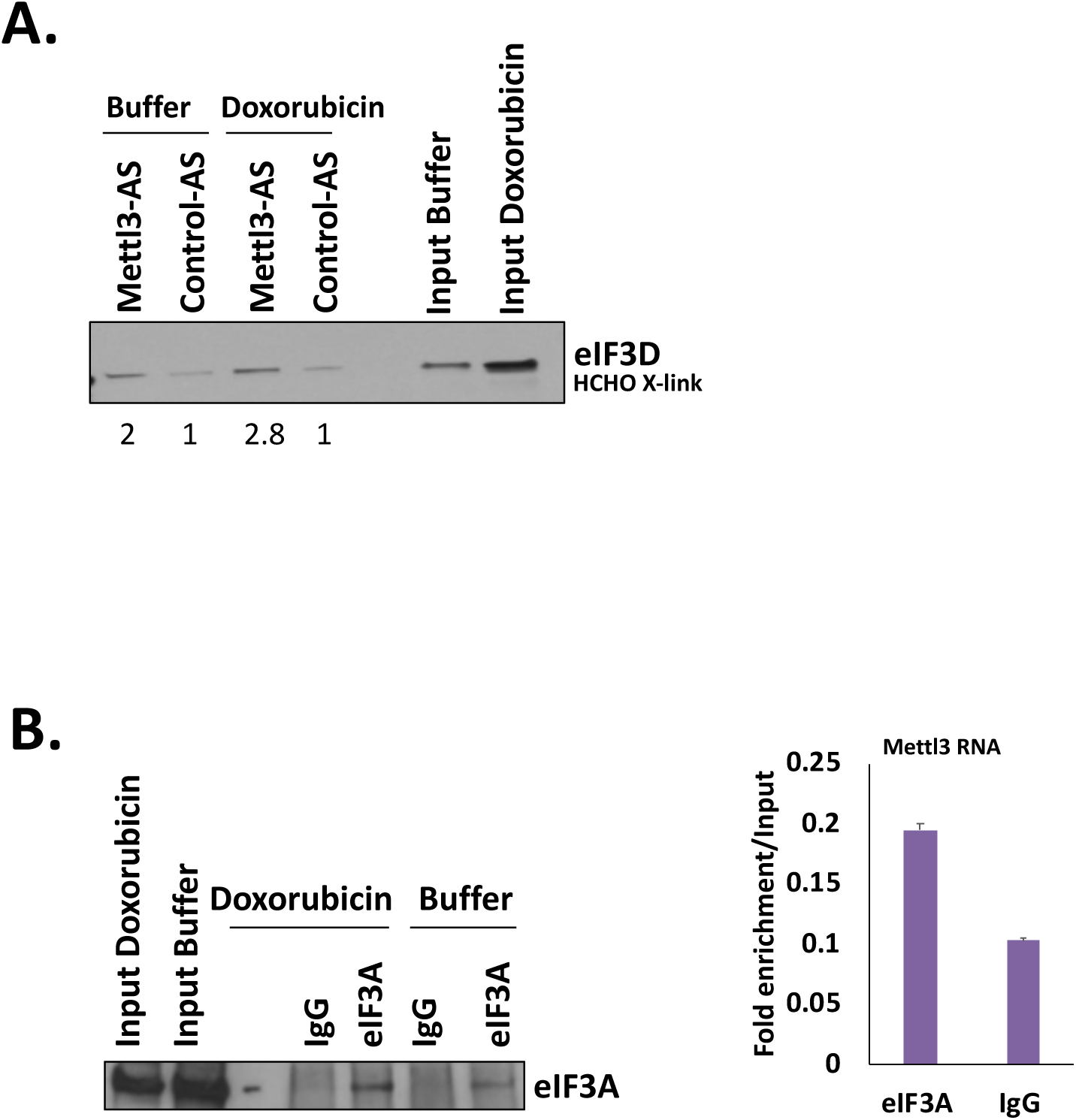
**A.** Western blot of eIF3d co-purification with biotin-tagged antisense to METTL3 mRNA after formaldehyde crosslinking of doxorubicin-treated and untreated MCF7 cells (250nM, 12 h). EIF3d quantification below normalized for Input levels and for METTL3 mRNA purified with antisense and quantified by qPCR normalized for Input RNA levels. **B.** Western blot of eIF3a immunoprecipitation that was analyzed by qPCR for co-purification of METTL3 mRNA (side graph) in 500nM doxorubicin-treated and untreated MCF7 cells, after in vivo UV crosslinking. Data are average of 3 replicates +/−SEM. See also Fig. 3, Tables S1, S2A-B.

**Fig. S4.**
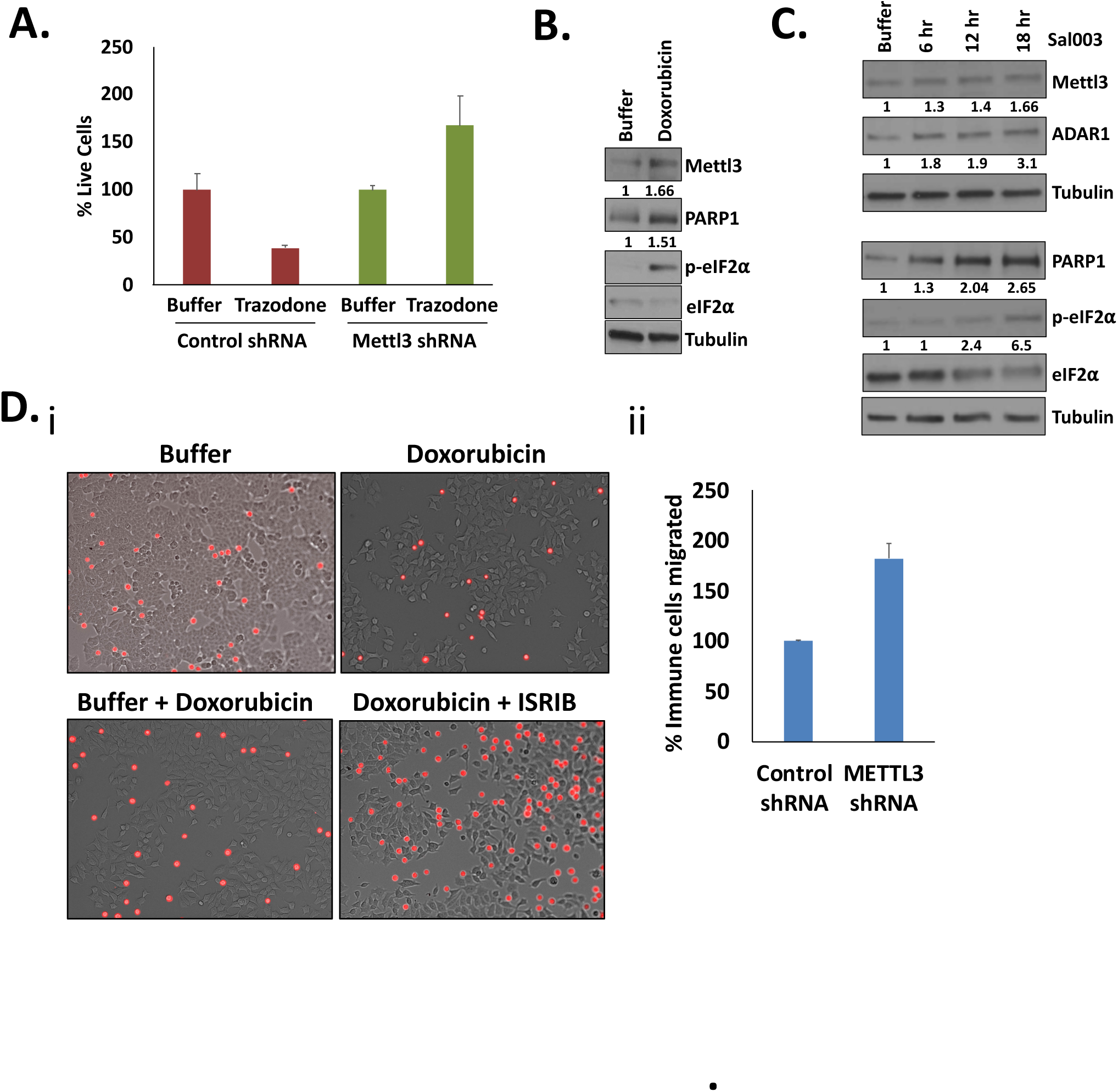
**A.** To test whether the reversal of chemosurvival by Trazodone (5uM) in Fig. 4Eii, was due to METTL3, doxorubicin (500nM) treated control shRNA MCF7 cells were compared with shMETTL3 cells (percentage chemosurviving cells) after treatment with buffer or Trazodone to override the eIF2α phosphorylation pathway and block METTL3-induced chemosurvival. Treatment of shMETTL3 cells with doxorubicin and with Trazodone that overrides eIF2α phosphorylation, does not cause additional loss of chemosensitivity, indicating that METTL3 and the eIF2α phosphorylation pathway targeted by this inhibitor, are in the same pathway and that the reduced chemosurvival with Trazodone in Fig. 4Eii is due in part to reduced METTL3. **B-C**. Western blot analysis of PARP, ADAR1, phospho-eIF2α, and METTL3 with 500nM doxorubicin or 10uM Sal003 treatment for 18hrs, compared to control buffer. **D. i.** Immune cell migration assay with 500nM doxorubicin treated MCF7 cells and with cells co-cultured with 1uM ISRIB for 18hrs. **ii.** Bar graph shows more immune cell migration upon METTL3 depletion. Data are average of 3 replicates +/−SEM. Actin and Tubulin are loading controls for Western blots. See also Fig. 4, Tables S3-5.

**Fig. S5.**
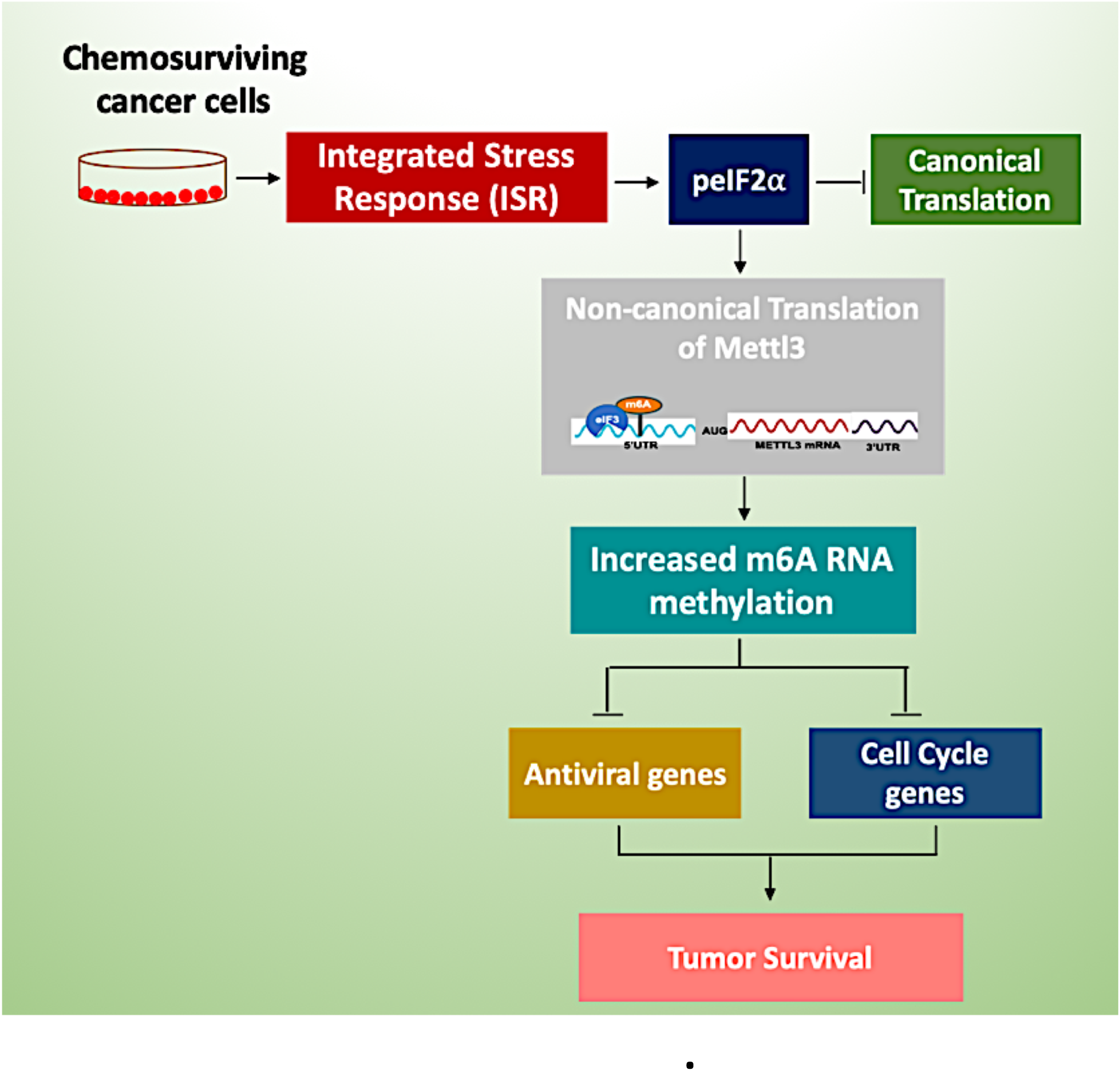
Model for therapy-induced ISR regulating tumor survival via METTL3 regulation.

## Supplemental Table Legends

**Table S1.** TMT spectrometry analysis of 500nM doxorubicin (1 day) treated versus untreated MCF7 cells. The cells were sonicated to disrupt nuclei and cells completely, prior to TMT spectrometry.

**Table S2. A.** Biotin-tagged METTL3 RNA affinity purification against biotin-tagged control scrambled N1 antisense purification in untreated versus 500nM doxorubicin (1 day) treated in vivo formaldehyde crosslinked MCF7 cells, followed by TMT spectrometry analysis. **B.** m6A IP in 500nM doxorubicin (1day) treated versus untreated MCF7 cells versus control IgG antibody, followed by TMT spectrometry analysis.

**Table S3.** Microarray analysis of RNA levels affected by METTL3 depletion and m6A immunoprecipitated (IP) RNAs.

**Table S4.** TMT spectrometry analysis of METTL3 depletion compared to control shRNA in 500nM doxorubicin treated and untreated MCF7 cells, and of 500nM doxorubicin (1 day) treated versus untreated MCF7 cells.

**Table S5.** Commonly regulated genes from datasets in Tables S3-S4 of M6A IP RNAs in doxorubicin-treated MCF7 cells, which are regulated at the RNA or protein level upon METTL3 depletion: M6A IP RNAs in doxorubicin-treated cells versus untreated MCF7 cells, which are also regulated in RNA levels or proteome in shMETTL3 cells versus control shRNA cells with doxorubicin treatment, reveal that m6A decreases cell cycle RNAs, anti-viral immune RNAs, metabolism RNAs (up in m6A IP and up in shMettl3 compared to control shRNA doxorubicin treated cells), and promotes protein levels of invasion genes (up in m6A IP and down in shMettl3 compared to control shRNA doxorubicin treated cells), as well as m6A IP targets, PARP, and ADAR1.

**Table S6.** Primers and antibodies used with their sources.

